# Cytoplasmic Microtubule Organizing Centers Regulate Meiotic Spindle Positioning in Mouse Oocyte

**DOI:** 10.1101/2020.06.25.172684

**Authors:** Daniela Londono Vasquez, Katherine Rodriguez-Lukey, Susanta K. Behura, Ahmed Z. Balboula

**Affiliations:** Animal Sciences Research Center, University of Missouri, Columbia, MO 65211, USA; University of Cambridge, Department of Genetics, Downing Street, Cambridge, CB2 3EH, UK

**Keywords:** MTOC, Microtubule, Oocyte, Meiosis, Spindle positioning

## Abstract

During oocyte meiosis, migration of the spindle and its positioning must be tightly regulated to ensure elimination of the polar bodies and provide developmentally competent euploid eggs. Although the role of F-actin in regulating these critical processes has been studied extensively, little is known whether microtubules (MTs) participate in regulating these processes. Here, we characterize a pool of MTOCs in the oocyte that does not contribute to spindle assembly but instead remains free in the cytoplasm during metaphase I (metaphase cytoplasmic MTOCs; mcMTOCs). In contrast to spindle pole MTOCs, which primarily originate from the perinuclear region in prophase I, the mcMTOCs are found near the cortex of the oocyte. At nuclear envelope breakdown, they exhibit robust nucleation of MTs, which diminishes during polar body extrusion before returning robustly during metaphase II. The asymmetric positioning of the mcMTOCs provides the spindle with a MT-based anchor line to the cortex opposite the site of polar body extrusion. Depletion of mcMTOCs, by laser ablation, or manipulating their numbers, through autophagy inhibition, revealed that the mcMTOCs are required to regulate the timely migration and positioning of the spindle in meiosis. We discuss how forces exerted by F-actin in mediating movement of the spindle to the oocyte cortex are balanced by MT-mediated forces from the mcMTOCs to ensure spindle positioning and timely spindle migration.

## INTRODUCTION

Mammalian oocytes enter meiosis during early fetal life. Soon after birth, meiotic oocytes undergo a lengthy arrest at the dictyate stage of the prophase I of the first meiotic division (MI) ^1^. At the age of puberty, gonadotropin cues allow prophase I-arrested oocytes to resume MI evident by breakdown of the nuclear envelope (NEBD) and formation of a central bipolar spindle ^2-4^. The central positioning of the spindle is required to establish proper kinetochore-microtubule (MT) attachments and to protect against aneuploidy ^5^. The position of the spindle dictates the plane of cell division ^6^ and therefore, in contrast to mitotic cells where a centrally positioned spindle allows symmetrical cell division, the meiotic spindle must migrate towards the cortex for the highly asymmetric meiotic divisions ^6,7^. Such peripheral positioning of the spindle is critical to extrude the tiny polar body (PB) thereby retaining the great majority of the cytoplasm containing maternal RNAs and proteins for the egg to support early embryonic development ^8,9^. It is essential to understand the critical events of spindle positioning and migration, which are required for the fidelity of chromosome transmission to the next generation.

To date, F-actin and its regulatory molecules represent the only cytoskeletal components known to regulate spindle migration and positioning in the mammalian oocyte. It has been shown that perturbation of F-actin, but not of MTs, impairs spindle migration in oocytes and that the resulting increased symmetry of cell division results in infertility ^7,10-12^. This contrasts with mitotically dividing somatic cells, where positioning of the spindle at the center of the cell is primarily regulated by the interaction of the astral MTs originated from its poles with the cell cortex ^13^. Such astral MTs are nucleated by centrosomes, centriole pairs surrounded by peri-centriolar material (PCM). By contrast, mammalian oocytes lack classic centrosomes ^14^ because centrioles are lost during early oogenesis through an unknown mechanism. The numerous acentriolar MT organizing centers (MTOCs) of the oocyte ^14,15^ are still able to nucleate astral-like MTs but these are short and unable to extend to the cortex when the spindle is centrally positioned ^15-17^. These observations have enforced the notion that MTs have no role in regulating central spindle positioning and migration. It is therefore surprising that when mouse oocytes are treated with nocodazole, a MT depolymerizing agent, chromosomes migrate towards the cortex at a higher speed and at an earlier time than in control oocytes ^12,18^. These unexplained observations suggest that MTs have, yet unknown, role(s) in regulating spindle migration and positioning in mammalian oocytes.

Based on MTOC behavior, three different classes of MTOCs were characterized in meiotic oocytes. In prophase I-arrested oocytes, the MTOCs (class I) are initially found in the perinuclear region ^14^. Then, before NEBD, these perinuclear MTOCs undergo distinct processes of decondensation, stretching and redistribution into a large number of smaller MTOCs^19,20^. The fragmented MTOCs are then clustered and sorted to form two poles necessary to assemble a bipolar spindle ^15,21,22^. Another pool of MTOCs is also present in the cytoplasm during NEBD. Some of these cytoplasmic MTOCs (class II) migrate from the periphery to the center of the oocyte, where they participate in spindle pole formation (polar MTOCs, pMTOCs; classes I and II) ^15,19^. Another subset of cytoplasmic MTOCs (class III), hereafter referred to as metaphase cytoplasmic MTOCs (mcMTOCs) does not contribute to spindle formation and has yet unknown biological significance for oocyte meiosis.

Here, we show that the pMTOCs and the mcMTOCs of MI oocytes represent two different functional sets. We find by 3D time-lapse imaging that mcMTOCs localize asymmetrically opposite the site of F-actin enrichment where the first PB is extruded. Super-resolution Stimulated Emission Depletion (STED) microscopy reveals that mcMTOCs are able to nucleate MTs that connect the spindle to the cortex. Importantly, we show that by increasing mcMTOC numbers following treatment with an inhibitor of autophagy or by depleting them by laser ablation, the meiotic spindle becomes abnormally positioned leading to aneuploidy. Our results suggest a model whereby the role of F-actin in mediating movement of the meiotic spindle to the cortex is balanced by forces exerted from mcMTOCs to ensure the timely migration and accurate positioning of the spindle in the oocyte.

## RESULTS

### MTOCs in the cytoplasm and at the spindle poles of meiotic oocytes

The multiple MTOCs of prophase I-arrested oocytes follow distinct patterns of behavior. Immediately before NEBD, perinuclear MTOCs become fragmented in three phases; decondensation, stretching and repositioning towards the spindle poles through the sequential actions of Polo-like kinase 1, BicD2-anchored dynein, and the KIF11 motor protein ^19,20^. Soon after NEBD, a second group of cytoplasmic MTOCs migrates from the periphery to the center of the oocyte, where it participates with perinuclear MTOCs in spindle assembly ^15,19^. Finally, a third population of MTOCs persists in the cytoplasm of MI oocytes at metaphase (here termed mcMTOCs) ^17,23-26^, whose role in meiosis is unknown. To follow the behavior of mcMTOCs during MI, we employed live imaging using 3D confocal microscopy to visualize prophase I-arrested oocytes (collected from CF-1 mice) expressing GFP-tagged Aurora A Kinase (AURKA-GFP, an integral component of MTOCs in mouse oocytes ^27-29^ and H2B-mCherry (H2B-mCh) to label MTOCs and DNA, respectively. As previously reported, perinuclear MTOCs became fragmented into small multiple MTOCs at NEBD before they sorted and re-clustered at the two spindle poles as pMTOCs (Fig. 1A; Supplementary Movie 1). We also observed cytoplasmic MTOCs in the cytoplasm at NEBD, some of which migrated towards the oocyte center to contribute to spindle formation as pMTOCs, whereas others, mcMTOCs, remained free in the cytoplasm during metaphase I (Met I) after the bipolar spindle had formed (Fig. 1A; Supplementary Movie 1). We carried out 3D reconstruction of entire oocytes to examine the number (Fig. 1 B,C) and volume (Fig. D, E) of the mcMTOCs (Supplementary Movie 2). In contrast to pMTOCs which undergo a time-dependent decrease in number and increase in volume (due to MTOC clustering) as the oocyte proceeds to Met I, the mcMTOCs displayed the opposite pattern and showed a time-dependent increase in number and volume during pro-Met I and Met I (Fig. 1A,B,D; Supplementary Movies 1 and 2). Each oocyte had a variable number of mcMTOCs (between 4 and 12) at Met I located on different focal planes (Supplementary Movie 3). The mcMTOCs became less distinct, appearing to be decreased in number and volume during anaphase I (Ana I) and telophase I (Telo I) before regaining their metaphase appearance as the oocytes arrested in Met II (Fig. 1A-E; Supplementary Movies 1 and 2). We confirmed that these AURKA-positive foci were indeed MTOCs by showing the colocalization of pericentrin, another integral component of PCM in meiotic oocytes ^30,31^ (Supplementary Fig. 1A). To confirm our observations, we fixed CF-1 oocytes at different developmental stages (Prophase I, Met I, Ana I/Telo I and Met II) and immunostained them to reveal the MTOC markers, γ-tubulin, pericentrin and CEP192^19,31-33^. This also revealed pMTOCs and mcMTOCs at Met I (Fig. 1F, G) that were associated with MT asters (α-tubulin staining, Fig. 1F). Similar findings were observed in Met I oocytes from C57BL/6 mice (Supplementary Fig. 1B). Just as we observed in time-lapse imaging of MTOCs in living oocytes, the mcMTOCs became less distinct in Ana I/Telo I of fixed oocytes (Fig. 1F).

**Figure 1:**
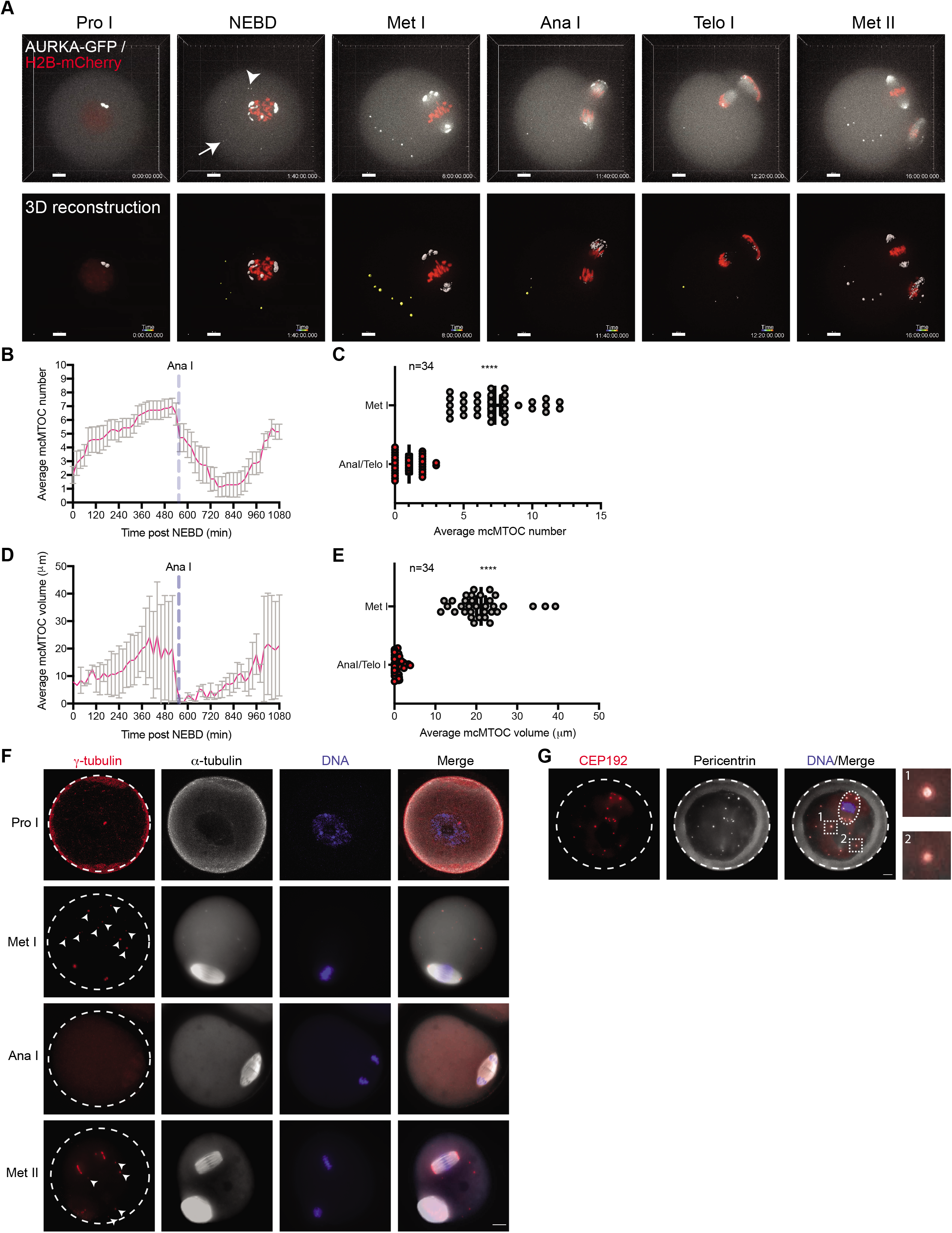
Two different sets of MTOCs are present in acentriolar oocytes. (A) Time-lapse confocal microscopy of MTOCs in live oocytes. Full-grown prophase-I oocytes were injected with cRNAs encoding *H2b-mCherry* (red) and *Aurka-Gfp* (pseudo white), incubated in milrinone-containing CZB medium for 3 h, followed by *in vitro* maturation. Shown are representative images (Z-projection of 16 sections every 3□μm) from a time course (see Supplemental Movie 1). Lower panels represent a 3D reconstruction of MTOCs from oocyte shown in upper panels (see Supplemental Movie 2). Fluorescence images were captured every 20 min. The scale bar represents 10□μm. The white arrowhead represents cytoplasmic MTOC that participate in spindle pole formation as a pMTOC. The white arrow represents cytoplasmic MTOC that remains in the cytoplasm during Met I (mcMTOC). (B,D) Quantification of average MTOC number and MTOC volume over time during meiosis I, respectively. Error bars show S.D. Dashed blue lines represent the time of Anaphase I onset. (C,E). Quantification of average MTOC number and MTOC volume, respectively, during metaphase I (Met I) and anaphase I/telophase I (Ana I/Telo I) stages. The data are expressed as mean ± SEM. Student t-test was used to analyze the data. Values with asterisks vary significantly, ****P < 0.0001. The total number of analyzed oocytes (from three independent replicates) is specified above each graph. (F) Fully grown prophase-I-arrested oocytes were *in vitro* matured for 0 h (prophase I, Pro I), 8 h (Met I), 9 h (Ana I/Telo I) or 16 h (Met II) prior to fixation and immunocytochemistry using γ-tubulin and α-tubulin antibodies to label MTOCs (red) and microtubules (pseudo grey). DAPI was used to detect DNA (blue). (G) Fully grown prophase-I-arrested oocytes were *in vitro* matured for 8 h (Met I) followed by fixation and immunostaining using CEP192 (red) and pericentrin (grey) antibodies to label MTOCs. DAPI was used to detect DNA (blue). Shown are representative confocal Z-projections. Arrowheads represent mcMTOCs. A total of 184 oocytes were examined. Scale bars represent 10□μm.

### mcMTOCs undergo three patterns of directional movement during MI

To dissect the directional motion and kinetics of the mcMTOCs, we tracked their movement using 3D reconstructions over time in pro-Met I oocytes expressing AURKA-GFP. In contrast to pMTOCs which primarily originate from perinuclear MTOCs at NEBD, the mcMTOCs formed at the periphery of the oocyte (Fig. 2A; Supplementary Movie 4 and 5) and then appeared to undergo three phases of directional movement. In the first phase, from NEBD to early Met I, the peripheral mcMTOCs moved towards the oocyte’s center with an average speed of 0.09 ± 0.007 μm min^− 1^ and a maximum speed of 0.18 ± 0.005 μm min^− 1^. During this phase, the average volume of the mcMTOCs increased, in some cases due to mcMTOCs merging with each other (Fig. 2A; Supplementary Movie 4 and 5). The second was marked by the slowing of mcMTOC movement to an average speed of 0.06 ± 0.009 μm min^− 1^ allowing them to remain in confined areas of the cytoplasm (Fig. 2B; Supplementary Movie 4). During this phase, the pMTOCs underwent active clustering whereas the mcMTOCs remained apart. The third phase occurred during Ana I and Telo I when the mcMTOCs showed a reversal of their behavior in phase I; they displayed a drastic reduction in volume and migrated towards the cortex with an average speed of 0.12 ± 0.01 μm min^− 1^ and a maximum speed of 0.22 μm min^− 1^ (Fig. 2C; Supplementary Movies 4 and 6). In as many as 50% of oocytes (14/28), we were not able to observe mcMTOCs during Ana I/Telo I but in all cases when they could be observed, the mcMTOCs migrated towards the region of cortex opposite direction the site of extrusion of the first PB. The mcMTOCs showed independent directional movement to cytoplasmic droplets and so do not reflect overall cytoplasmic movements (Supplementary Fig. 2). To exclude the possibility that the decrease in mcMTOC size during Ana I/Telo I is due to AURKA protein degradation, we generated a *Cep192-eGfp* reporter CF-1 mouse model (Supplementary Fig. 3A) to allow for the precise tracking of mcMTOCs during MI. Briefly, a construct harboring the EGFP reporter gene was integrated into the mouse genome via CRISPR/Cas9-mediated homology-directed repair. The EGFP reporter was fused at the C-terminus of the endogenous mouse *Cep192*. Thus, in this model, CEP192 is just labeled, not perturbed or overexpressed. Accordingly, *Cep192-eGfp* oocytes expressed similar levels of CEP192 and are phenotypically normal to WT oocytes in terms of meiotic progression (Supplementary Fig. 3B), chromosome alignment, spindle morphology, positioning, and migration (data not shown). Using immunocytochemistry, we confirmed that endogenous CEP192-eGFP colocalizes with all pericentrin-positive MTOCs (Supplementary Fig. 3C). Importantly, CEP192-eGFP oocytes showed similar mcMTOC behavior during MI where variable number of mcMTOCs were scattered in the cytoplasm during Met I (Supplementary Fig. 3D). During Ana I/Telo I, similar to AURKA-labeled mcMTOCs, endogenously labeled mcMTOCs (CEP192-eGFP) underwent a decrease in size and number while migrating towards the cortex (Supplementary Fig. 3D; Supplementary Movie 7). Consistently, we were not able to observe mcMTOCs during Ana I/Telo I in ∼ 46% of oocytes (35/76) prior to their reappearance again at Met II. Taken together, our observations confirm that meiotic oocytes have two different pools of MTOCs during Met I and suggest that mcMTOCs differ from pMTOCs in their function.

**Figure 2:**
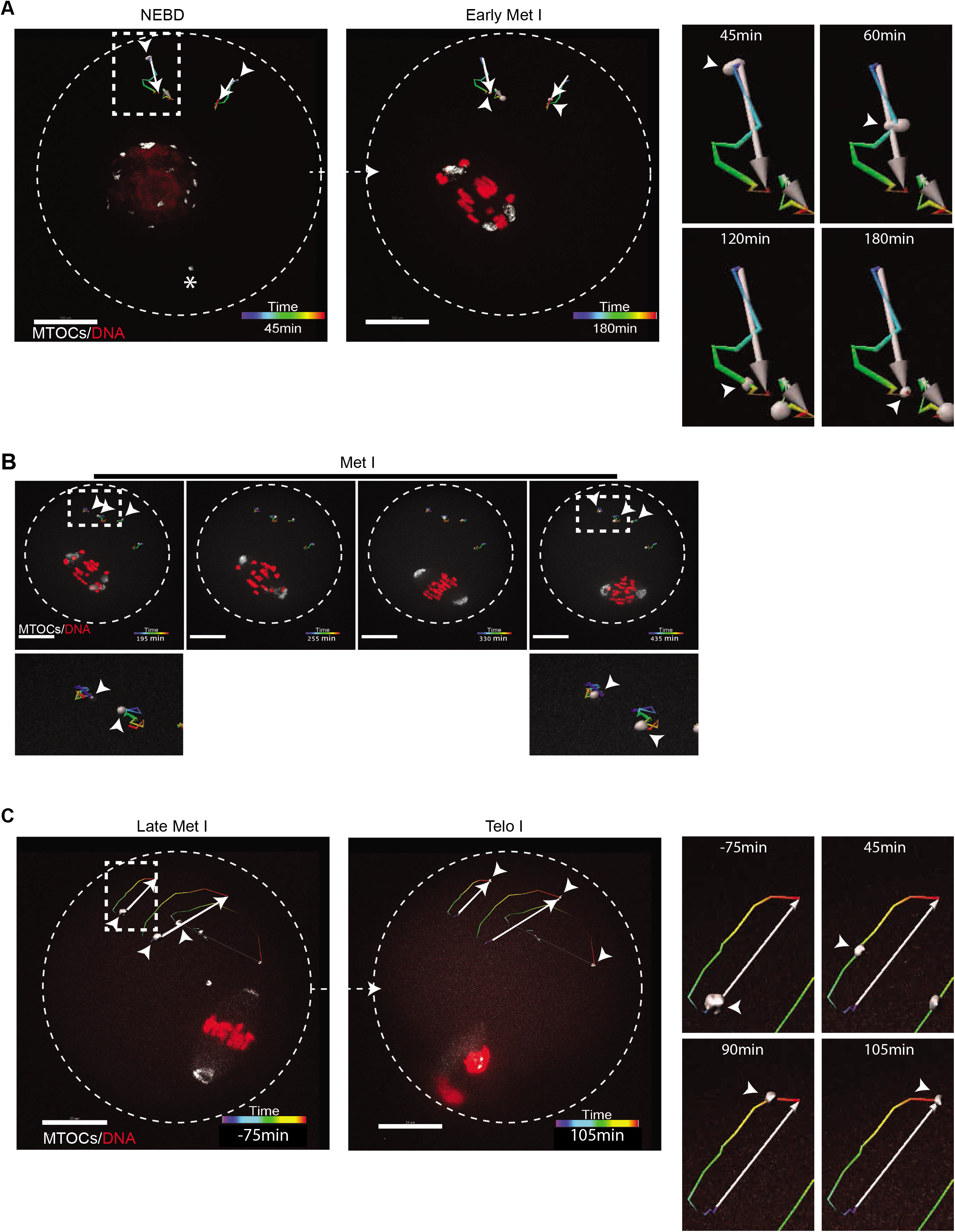
mcMTOCs exhibit three different directional behaviors during meiosis I. Tracking of 3D reconstructed mcMTOCs from time-lapse confocal microscopy in live oocytes during three different phases: (A) nuclear envelope breakdown (NEBD)-early metaphase I (Met I, see Supplemental Movie 4 and 5), (B) Met I (see Supplemental Movie 4) and (C) late Met I-telophase I (Telo I), see Supplemental Movie 5 and 6. Full-grown prophase-I oocytes were injected with cRNAs encoding *H2b-mCherry* (red) and *Aurka-Gfp* (pseudo white), incubated in milrinone-containing CZB medium for 3 h prior to *in vitro* maturation. Shown are representative images of 3D reconstructed Z-projection of 16 sections every 3□μm from a time course. Fluorescence images were captured every 15 min (time, h:min). The scale bar represents 20□μm. Right panels show a zoomed area of mcMTOC tracking over time. White arrows represent the overall direction of displacement calculated automatically by Bitplane Imaris software. Arrowheads represent mcMTOCs. The total number of analyzed oocytes is 60.

### mcMTOCs localize asymmetrically to anchor the spindle to the cortex

To determine whether the mcMTOCs were physically connected to pMTOCs, we employed immunocytochemistry and STED super-resolution microscopy to visualize the MTOCs in relation to MTs and F-actin. We confirmed that F-actin formed a cage around the spindle as previously reported ^11,34^ but could not detect any direct connection of F-actin between the pMTOCs and the mcMTOCs (Supplementary Fig. 4A). In contrast, we detected MTs originating from the mcMTOCs and linking them to both the oocyte cortex and the pMTOCs and spindle (Fig. 3A,B). In addition, we also observed MTs connecting mcMTOCs with each other. Relatively short astral-like MTs could not reach the oocyte cortex unless they bind mcyMTOCs (Fig. 3A,B). Thus, mcMTOCs enable MTs to bridge the gap between the spindle and the cortex. It was also evident from our observations that the mcMTOCs were asymmetrically distributed in the oocyte cytoplasm leading us to consider their relationship in time and space to the positioning of the spindle itself. In prophase arrest oocytes, the nucleus (Germinal vesicle, GV) is usually found in a centralized location ^35-38^. Consequently, the spindle also forms around the chromatin at or near the center of the oocyte before it migrates towards the cortex during late bMet I to allow asymmetrical cell division. In all the oocytes examined, we found that the mcMTOCs were all asymmetrically positioned from the GV stage and throughout NEBD and early pro-Met I when the spindle is still localized centrally (Fig. 3C; Supplementary Movie 8). Strikingly, when the spindle began its actin-mediated migration, this took place towards the opposite side of the oocyte to that occupied by mcMTOCs in the majority of the oocytes (38 out of 40 examined, Fig. 3 C-E). These findings suggest a model whereby mcMTOCs localize asymmetrically to anchor the spindle to the cortex in such a way as to oppose the F-actin mediated force that builds to direct the spindle to the cortex on the opposite face of the oocyte for PB extrusion (Fig. 7). This would account for the unexplained finding that nocodazole-induced MT depolymerization during MI results in the earlier migration of chromosomes towards the cortex at a relatively higher speed than in control oocytes (Supplementary Fig. 4B,C) and others ^12,18^. Thus, the mcMTOCs appear to enable opposing MT forces that counter spindle/chromosome migration that are necessary to regulate its timing and the final position of the spindle.

**Figure 3:**
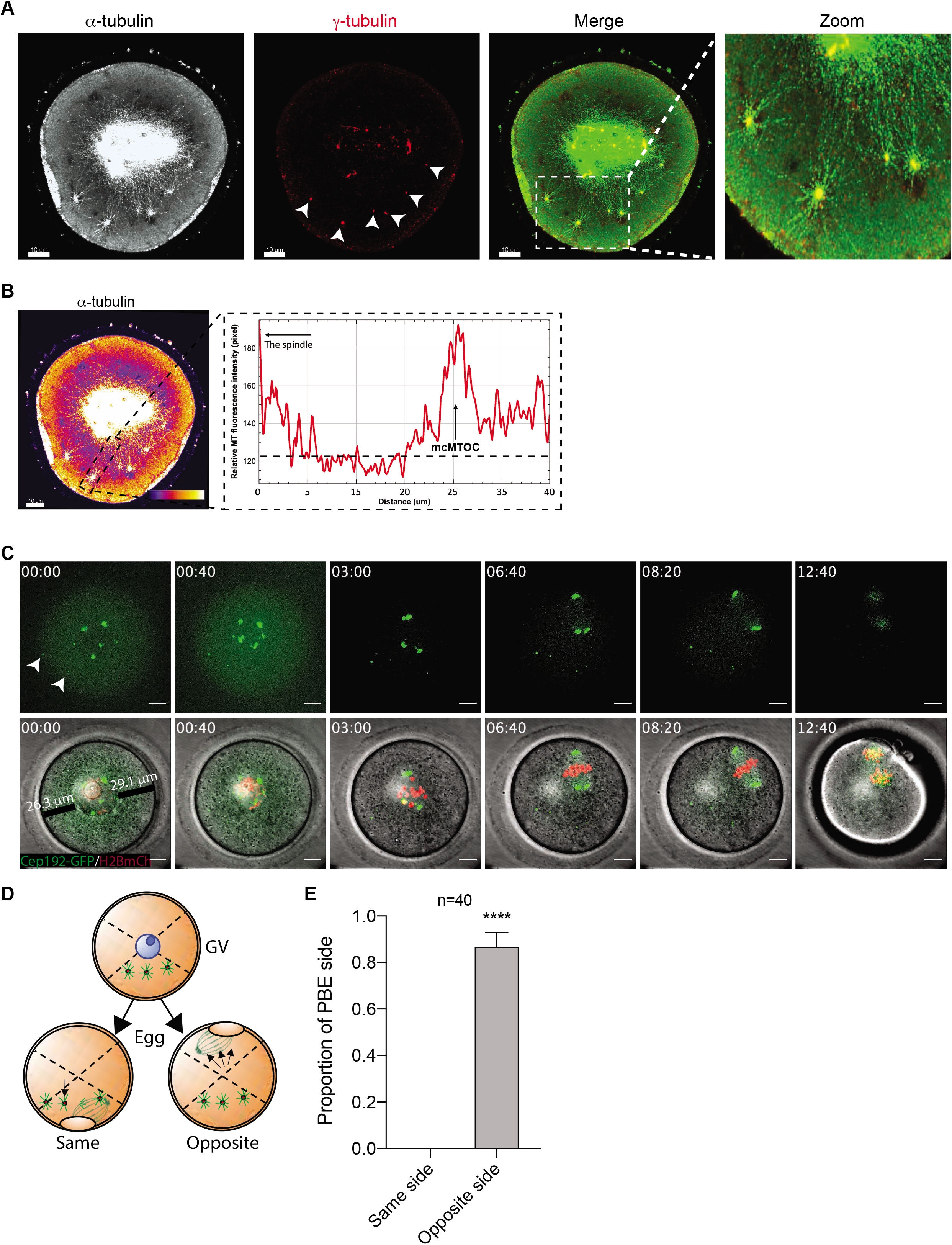
mcMTOCs anchor the spindle to the cortex. (A) Fully grown prophase-I-arrested oocytes were *in vitro* matured for 6 h (metaphase I) prior to fixation and immunocytochemistry using γ-tubulin and α-tubulin antibodies to label MTOCs (red) and microtubules (pseudo grey). Hoechst was used to detect DNA (blue). Fluorescence signals were detected under a 63X objective using STED super-resolution system. Shown is a representative image (Z-projection of 65 sections every 0.5□μm). (B) Example of fluorescence intensity of microtubules connecting pMTOCs, mcMTOCs and the cortex, using the ‘plot profiles’ function in ImageJ. The scale bar represents 10□μm. (C) Representative images (Z-projection of 16 sections every 3□μm) of time-lapse confocal microscopy of a live oocyte expressing AURKA-GFP (MTOCs) and H2B-mCherry (chromosomes) from a time course (see Supplemental Movie 8). Fluorescence and bright-field images (lower panels) were captured every 20 min (time, h:min). Arrowheads represent mcMTOCs. Scale bars represent 10□μm. (D) Schematic diagram shows how the proportion of polar body extrusion side in relation to mcMTOC position was assessed. (E) Quantification of the proportion of polar body extrusion side from “C” according to “D”. The data are expressed as mean ± SEM. Student t-test was used to analyze the data. Values with asterisks vary significantly, ****P < 0.0001. The total number of analyzed oocytes (from three independent replicates) is specified above the graph.

### mcMTOCs regulate spindle positioning in acentriolar oocytes

To determine the function(s) of mcMTOCs during MI and to test the above hypothesis, we selectively depleted mcMTOCs by two-photon laser ablation (Fig. 4A). The two-photon laser microscope has the advantage of offering deeper tissue penetration enabling efficient ablation and minimizing off-target effects ^39^. We first microinjected prophase I-arrested oocytes with cRNAs encoding *Aurka-Gfp* and *eGFP-Eb3* to label MTOCs and MTs, respectively. We then marked small cuboidal regions surrounding each mcMTOC, which we then exposed to a 925 nm wavelength laser to ablate the mcMTOCs, (Fig. 4B; Supplementary Movie 9). We ensure reduction of fluorescence of each mcMTOC to background levels (compare images before ablation, Fig. 4B, upper panels, to after ablation, Fig. 4B, lower panels) before ablating the next. Importantly, laser ablation not only depleted the mcMTOCs but also disrupted their associated nucleation of MTs (Fig. 4C). We also exposed control oocytes to the same protocol by ablating random areas of the cytoplasm (same size and numbers of mcMTOCs) adjacent to but not overlapping with the mcMTOCs. We confirmed the efficiency of mcMTOC depletion by immunostaining a subset of oocytes to reveal γ-tubulin and were only able to detect γ-tubulin foci at spindle poles but not in the cytoplasm (Supplementary Fig. 5). To assess spindle positioning, we arrested meiotic progression in Met I by incubating oocytes in meiotic maturation medium containing a proteasome inhibitor (MG-132). In this way, we could compare spindle positioning in such arrested oocytes for a longer time (Fig. 4A,D). This revealed that whereas in control Met I-arrested oocytes, the spindle maintained its position over a period of 9 h, the position of the spindle in mcMTOC-depleted oocytes was not stable and displayed abnormal movement. In contrast to control oocytes, the spindle in mcMTOC-depleted oocytes showed multiple migrations towards different cortical sides (Fig. 4D; Supplementary Movie 10 and 11). Consequently, the total distance traveled by the spindle and average spindle velocity (during 9 h time period) were significantly greater in mcMTOC-depleted oocytes than those in control oocytes (Fig. 4E-F). Following the onset of spindle movement in mcMTOC-depleted oocytes, the spindle poles lost their integrity in contrast to control oocytes, likely due to the imbalance of external and internal forces on spindle poles (Supplementary Movie 11). To assess the effect of mcMTOC depletion on meiotic progression, control and mcMTOC-depleted Met I oocytes were *in vitro* matured in MG-132-free medium. Depletion of mcMTOCs delayed PB extrusion (data not shown) and resulted in a significant increase in oocytes arrested at Met I compared to control oocytes (Fig. 4H). Because control-ablated oocytes extruded the first PB shortly after ablation (Fig. 4G; Supplementary Movie 12), statistical assessment of spindle positioning in mcMTOC-depleted oocytes was not feasible. However, similar to mcMTOC-depleted oocytes arrested with MG-132, we frequently observed spindle positioning defects and abnormal PB extrusion in *in vitro* matured mcMTOC-depleted oocytes (Fig. 4G; Supplementary Movie 13 and 14). Using an *in situ* chromosome counting technique^40,41^, we examined the karyotype of mcMTOC-depleted oocytes that reached Met II (eggs). A relatively higher proportion of mcMTOC-depleted eggs were aneuploid (18 out of 33 examined eggs) compared to controls (7 out of 32 examined eggs, Fig. 4I). Together this suggests that the mcMTOCs are required to position the Met I spindle.

**Figure 4:**
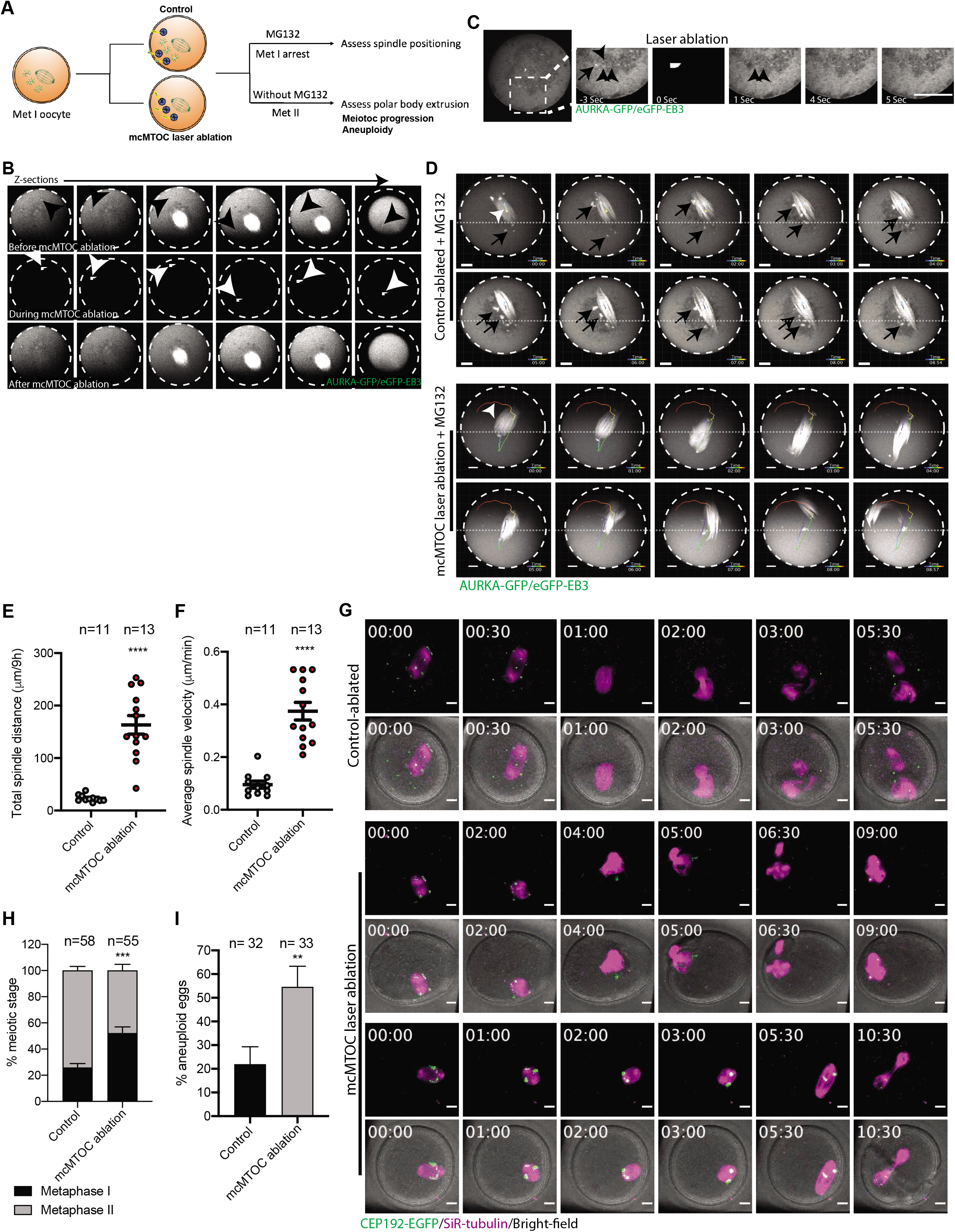
mcMTOCs are required to regulate spindle positioning in meiotic oocytes. (A) Schematic diagram shows the experimental design following mcMTOC depletion. (B-F) Oocytes expressing AURKA-GFP and eGFP-EB3 were *in vitro* maturated for 6 h (metaphase I, Met I), transferred to CZB medium with MG-132, followed by mcMTOC depletion using two-photon laser ablation. Small square area(s) surrounding mcMTOCs were marked and then exposed to a laser with 925 nm wavelength. Control oocytes were exposed to the same parameters except ablating random areas of the cytoplasm equal to the same size and number of mcMTOCs. (B) Z-sections of 3D time-lapse imaging of a live oocyte during mcMTOC ablation (see Supplemental Movie 9). Black arrowheads represent mcMTOCs before ablation (upper panels). White arrowheads represent laser beam targets (middle panels). (C) Z-sections of 3D time-lapse imaging of live oocyte to track microtubules (eGFP-EB3) following mcMTOC ablation. Black arrow represents an mcMTOC before ablation (−3 S). Black arrowheads track microtubules over time. The scale bar represents 20□μm. (D) Z-projection (16 sections every 3□μm) of 3D time-lapse microscopy of control-ablated (see Supplemental Movie 10) and mcMTOC-depleted (see Supplemental Movie 11) Met I oocytes while cultured in MG-132-containing medium for additional 9 h to track the spindle. Fluorescence images were captured every 3 min (time, h:min). White arrowheads represent the tracking path of all time periods. Black arrows represent examples of mcMTOCs on different focal planes. The scale bar represents 10□μm. (E) Quantification of total distance traveled by the spindle. (F) Quantification of average spindle velocity. The data are expressed as mean ± SEM. Student t-test was used to analyze the data. Values with asterisks vary significantly, ****P < 0.0001. (G) Control and mcMTOC-depleted *Cep192-eGfp* oocytes (MTOCs are labeled green) were *in vitro* matured in MG-132-free medium containing SiR-tubulin (to label microtubules, magenta) until Met II (16 h) and assessed for (H) meiotic progression and (I) aneuploidy. The data are expressed as mean ± SEM. Chi-square contingency test was used to analyze the data. Values with asterisks vary significantly, **P < 0.01, ***P < 0.001. The total number of analyzed oocytes in each group (from two independent replicates) is specified above each condition within each graph.

### Autophagy regulates mcMTOC numbers and spindle positioning in meiotic oocytes

In mitotic cells, autophagy plays an important role in regulating and maintaining the proper number of centrosomes where autophagy-deficient cells contained multiple centrosomes^42,43^. This led us to investigate the effects of inhibiting or inducing autophagy upon mcMTOC numbers and the consequences for meiosis. To this end, we chose to treat oocytes with 3-Methyladenine (3-MA), which inhibits autophagy by blocking autophagosome formation via the inhibition of type III Phosphatidylinositol 3-kinases (PI-3K). However, because 3-MA (autophagy inhibitor) blocks NEBD (data not shown), we treated oocytes with 3-MA immediately after NEBD. We also treated oocytes with rapamycin as an inducer of autophagy by adding the compound to the *in vitro* maturation medium during prophase I. In both cases, we allowed the treated oocytes (3-MA or rapamycin) together with controls to mature for 5 h post-NEBD prior to fixation and immunostaining using anti-CEP192 and anti-α-tubulin antibodies to label MTOCs and the spindle, respectively. We found that treatment with rapamycin resulted in a modest decrease in the number of mcMTOCs, compared to control oocytes (Fig. 5A-C). In contrast, treatment with 3-MA significantly increased (∼ 2-fold increase) the number of mcMTOCs, but not pMTOCs, compared to control oocytes (Fig. 5D-F). To determine the effect of such drug treatments upon spindle positioning, we used DIC imaging to track the position of chromosomes over time. Rapamycin-treated oocytes behaved in a similar way to controls; we could see no significant differences in the proportion of oocytes completing MI and extruding a PB (Fig. 6A,B). Moreover, rapamycin-treated oocytes showed no differences to controls in chromosome positioning (Fig. 6A; Supplementary Movie 15 and 16), in the total distance traveled by chromosomes until reaching the oocyte cortex; Fig. 6C), in the average time spent by chromosomes during migration to reach the cortex (Fig. 6D), or in the average speed of migrating chromosomes (Fig, 6E). By contrast, the increase in MTOC numbers following 3-MA treatment was associated with abnormal chromosome positioning and orientation with chromosomes moving in circles in oocytes treated with 3-MA (Fig. 6A; Supplementary Movie 17 and 18), likely due to frequent spindle misorientation (Fig. 6F; Supplementary Movie 20). Indeed, 3D reconstruction of the spindle and tracking the two spindle poles automatically showed abnormal spindle orientation and frequent spindle flipping (the phenotype that could not be observed in controls) during Met I in around 58% of 3-MA-treated oocytes (Fig. 6F; Supplementary Movie 19, 20 and 21). During MI, the spindle migrates along its longitudinal axis towards the cortex with minimal change in spindle orientation. Consistent with abnormal spindle positioning/orientation phenotype, and in contrast to DMSO-treated controls (Supplementary Movie 19), 3-MA-treated oocytes exhibited a significant increase in the rate of spindle orientation change during Met I (Fig. 6G,H; Supplementary Movie 20 and 21). Accordingly, we found a significant increase in the distance moved by chromosomes during Met I until reaching the closest point to the oocyte cortex (Fig. 6C). In line with our model that mcMTOC-mediated MTs anchor the spindle to the cortex opposite the PB extrusion side to position the spindle centrally, we found that increasing mcMTOC numbers by 3-MA treatment resulted in a significant increase in the duration of chromosome migration towards the cortex (Fig. 6D), resulting from their significantly (p < 0.001) reduced speed (Fig. 6E,I) in comparison to control oocytes. Indeed, we observed cases in which the chromosomes underwent segregation before the spindle had reached the cortex (Fig. 6J), resulting in enlarge PBs in around 21.05% of oocytes (Supplementary Movie 17, 18 and 20). The relatively weaker effect of rapamycin than 3-MA on spindle positioning could be attributed to our observation that rapamycin (at least at the used concentration) was not efficient to completely inhibit mcMTOCs and that mcMTOC numbers in rapamycin-treated oocytes remained within the range of mcMTOC numbers in the majority of control oocytes (between 4 and 12).

**Figure 5:**
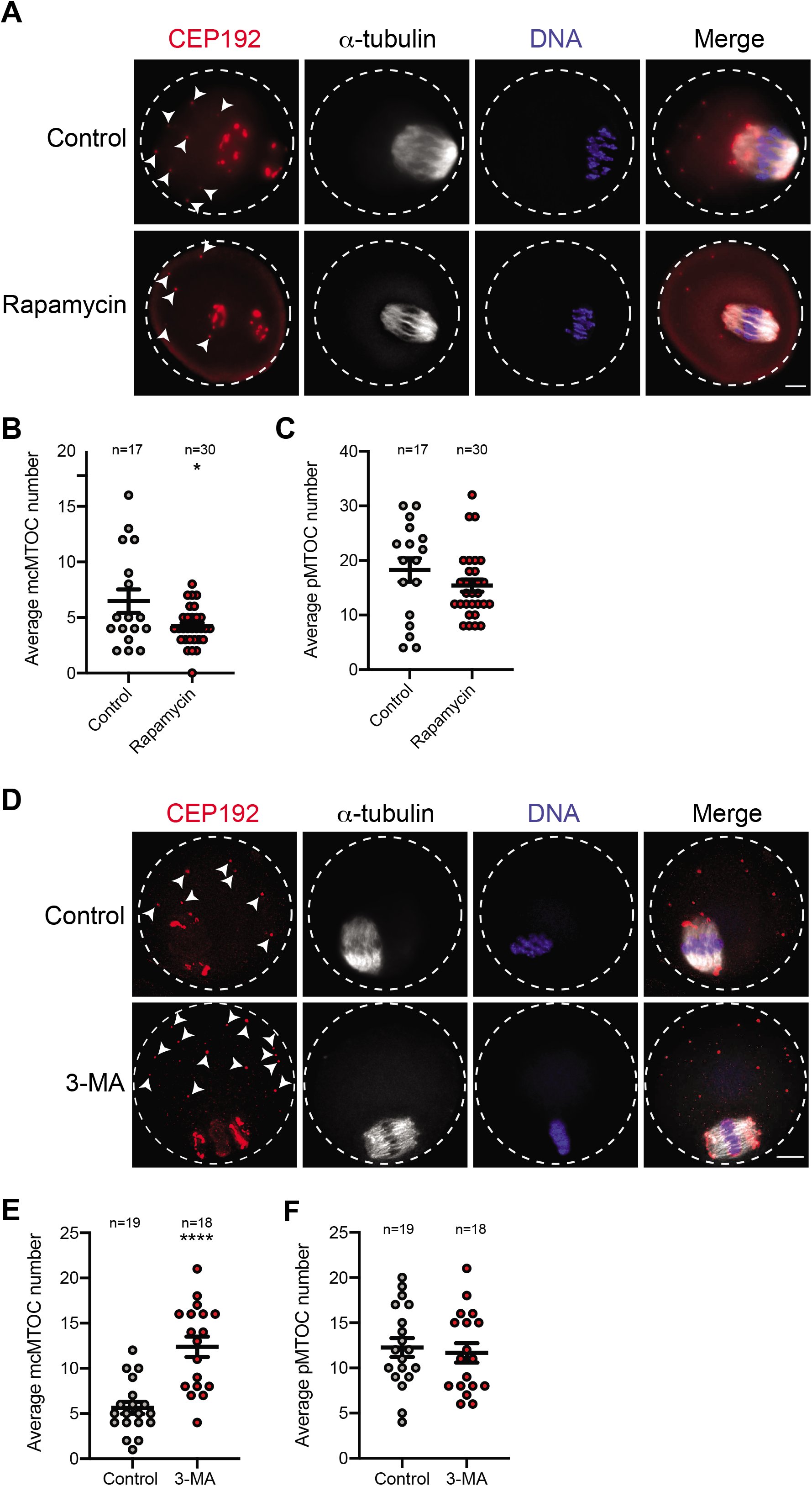
Autophagy regulates mcMTOC numbers in meiotic oocytes. (A) Full-grown prophase-I oocytes were divided into two groups and treated with DMSO or Rapamycin (added at 0 h after collection) followed by *in vitro* maturation until metaphase I (Met I, 7h). Met I oocytes were fixed and immunostained using CEP192 and α-tubulin antibodies to label MTOCs (red) and microtubules (pseudo grey). DAPI was used to detect DNA (blue). (B) Quantification of average mcMTOC numbers in “A”. (C) Quantification of average pMTOC numbers in “A”. (D) Full-grown prophase-I oocytes were divided into two groups and treated with DMSO or 3-MA (added at NEBD) followed by *in vitro* maturation until metaphase I (Met I, 7h). Met I oocytes were fixed and immunostained using CEP192 and α-tubulin antibodies to label MTOCs (red) and microtubules (pseudo grey). DAPI was used to detect DNA (blue). (E) Quantification of average mcMTOC numbers in “D”. (F) Quantification of average pMTOC numbers in “D”. Arrowheads represent mcMTOCs. The data are expressed as mean ± SEM. Student t-test was used to analyze the data. Values with asterisks vary significantly, *P < 0.05, ****P < 0.0001. The total number of analyzed oocytes (from two independent replicates) is specified above each graph.

**Figure 6:**
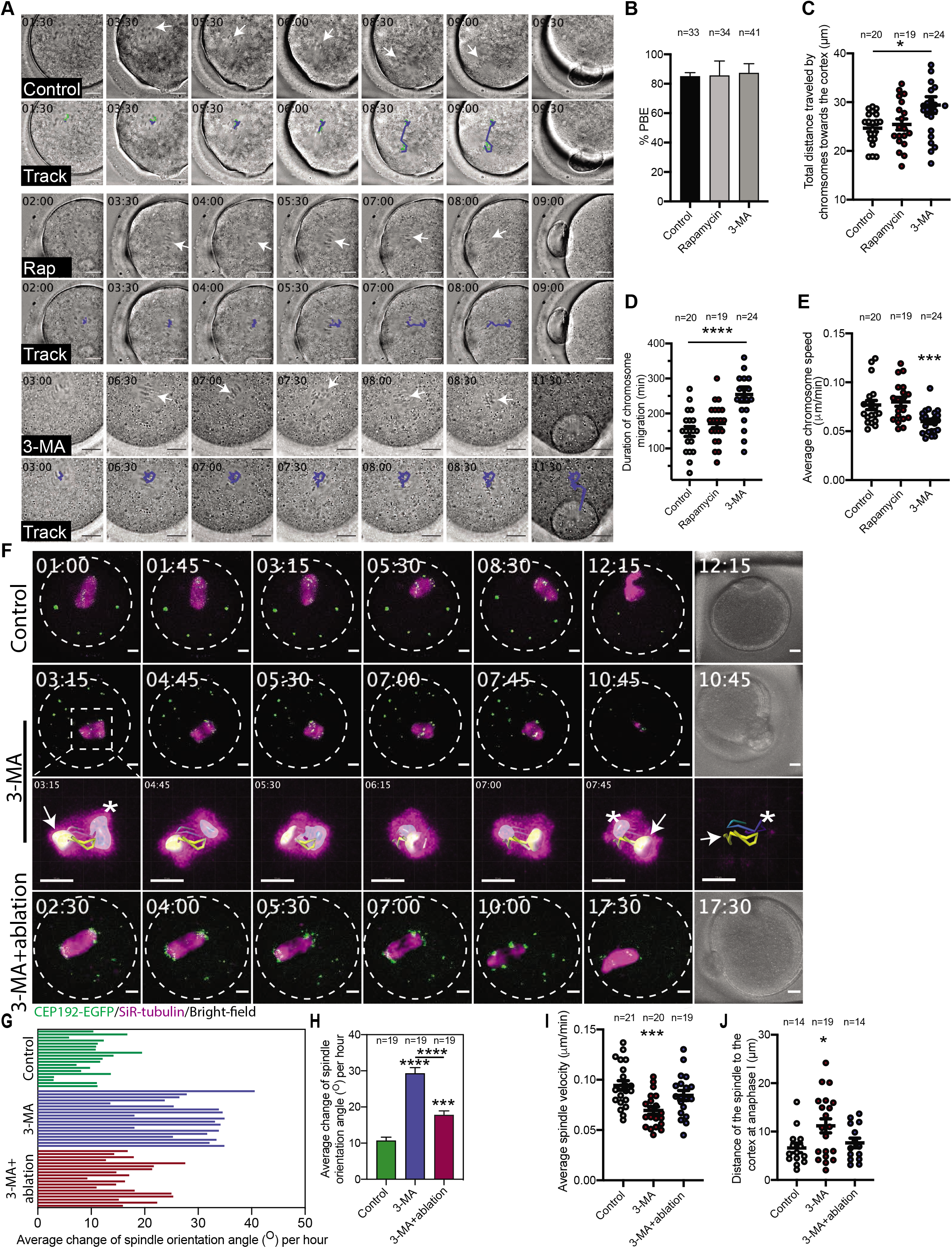
Autophagy regulates spindle positioning in meiotic oocytes. (A) Z-projection (16 sections every 3□μm) of time-lapse imaging of DMSO, Rapamycin (Rap) and 3-MA-treated oocytes (see Supplemental Movie 15, 16, 17 and 18, respectively). DIC images were captured every 30 min. White arrows represent the direction of chromosome displacement. Scale bars represent 10□μm. Lower panels of each group show the tracking path (blue) over time. (B) Quantification of the percentage of first polar body extrusion (PBE). (C) Quantification of the total distance traveled by chromosomes from the start of migration until reaching the nearest point to the cortex. (D) Quantification of the average time spent by chromosomes from the start of migration till reaching the nearest point to the cortex. (E) Quantification of average chromosome speed during migration until reaching the nearest point to the cortex. (F) Z-projection (13 sections every 5□μm) of time-lapse confocal imaging of *Cep192-eGfp* oocytes (MTOCs are labeled green) *in vitro* matured with SiR-tubulin (to label microtubules, magenta) and DMSO (control, upper panels), 3-MA (middle panels) or 3-MA+laser ablation (lower panels; 6 mcMTOCs were depleted, see Supplementary Fig. 6). Time-lapse imaging started 1 h post laser ablation. White arrows and asterisks indicate the two spindle poles. Scale bars represent 10□μm. (G) Quantification of the average change of spindle orientation angle per oocyte. Each bar represents an oocyte. (H) Quantification of the average change of spindle orientation angle per experimental group. (I) Quantification of average spindle velocity during migration until reaching the nearest point to the cortex. (J) Quantification of distance of the spindle to the nearest cortex at anaphase I. spindle velocity during migration until reaching the nearest point to the cortex. The data are expressed as mean ± SEM. One-way ANOVA was used to analyze the data. Values with asterisks vary significantly, *P < 0.05, ***P < 0.001, ****P < 0.0001. The total number of analyzed oocytes (from three independent replicates) is specified above each graph.

To confirm that spindle positioning defects in 3-MA-treated oocytes are due to increased mcMTOC numbers, we asked whether reducing mcMTOC numbers in 3-MA-treated oocytes can rescue, at least in part, spindle positioning defects. Using laser ablation, we depleted the “extra” mcMTOCs to achieve physiological mcMTOC numbers (6 to 8) in 3-MA-treated oocytes (Supplementary Fig. 6). Strikingly, ablating the “extra” mcMTOCs in 3-MA-treated oocytes partially rescued spindle positioning defects as evidenced by the decrease of oocytes exhibiting spindle flipping phenotype (4 out of 19 examined oocytes) when compared to 3-MA-treated oocytes (11 out of 20 examined oocytes, Fig. 6F; Supplementary Movie 20-22). Consistently, the average change of spindle orientation angle during Met I was greatly reduced when some of these extra mcMTOCs were ablated in 3-MA-treated oocytes (Fig. 6G,H). Because ablation of mcMTOCs (either all or some) causes a delay in PB extrusion (Fig. 4G, 6F and data not shown), compared to control-ablated oocytes, we could not assess the total distance traveled by the spindle during Met I. However, decreasing mcMTOC numbers in 3-MA-treated oocytes accelerated spindle migration towards the cortex and rescued the decrease in spindle velocity during migration (Fig. 6I). Importantly, decreasing mcMTOC numbers in 3-MA-treated oocytes by laser ablation greatly reduced the percentage of oocytes with enlarged PBs (5.26%), compared to non-ablated-3-MA oocytes (21.05%), resulting from the significant decrease in the distance between the spindle and the cortex at anaphase I (just before PB extrusion, Fig. 6J). Together, these data show that mcMTOC numbers must be regulated tightly during MI, and provide further evidence that mcMTOCs play an important role in regulating spindle positioning and its timely migration in mouse oocytes.

## DISCUSSION

To date, the only known function of acentriolar MTOCs in mouse oocytes is to assemble the spindle. Using 3D time-lapse confocal microscopy, we identify a subset of MTOCs that remain free in the cytoplasm during Met I of meiosis and which do not contribute to bipolar spindle assembly *per se*. In contrast to pMTOCs, which originate mainly from the perinuclear MTOCs in prophase I, the mcMTOCs originate exclusively from MTOCs present in the cytoplasm in prophase I. The mcMTOCs are first observed near the oocyte cortex at NEBD; they increase in number and size while moving to a central position during Met I. STED super-resolution microscopy revealed that MTs nucleated by the mcMTOCs connect one side of the spindle to the cortex during Met I. In Ana I/Telo I of MI, the mcMTOCs undergo a decrease in both number and size while migrating towards the cortex. When mcMTOCs functions were perturbed, either by laser ablation or treatment with 3-MA to inhibit autophagy and increase MTOC numbers, we have shown that the mcMTOCs play a role in regulating spindle positioning and the timing of its migration to the cortex.

To our knowledge, this is the first study of the function of mcMTOCs in living mammalian oocytes, which differ in several ways from the pMTOCs. The majority of pMTOCs, for example, originate from the perinuclear MTOCs, which never contribute to the mcMTOCs. Whereas the pMTOCs undergo a clustering-associated decrease in number and increase volume During pro-Met I/Met I, the mcMTOCs undergo a steady increase in both number and volume and rarely self-aggregate (∼1.5% of all examined mcMTOCs). Interestingly, inhibition of autophagy with 3-MA increased the number of mcMTOCs, but not pMTOCs. Finally, in contrast to pMTOCs, mcMTOCs participated in spindle positioning but never contributed to bipolar spindle assembly. Together, these observations suggest that mammalian oocytes have two different functional sets of MTOCs and raise the future important challenge to determine whether differences in their biochemical compositions underlie their differences in function.

The primary function of the spindle is to provide the machinery for faithful chromosome segregation. This is achieved in a series of critical, non-overlapping steps. First, during pro-Met I and early Met I, the spindle is assembled and positioned at or near the oocyte’s center. Second, during the late Met I, the spindle migrates towards a sub-cortical location to allow asymmetrical cell division. Third, the spindle rotates from a parallel to a perpendicular position in relation to the cortex to allow PB extrusion. Many studies have emphasized the roles of F-actin and its motor proteins in regulating spindle positioning and migration ^10-12^ and two models have been proposed to explain how F-actin regulates spindle positioning and migration. In the first, F-actin enrichment at the cortex provides a spindle pulling force ^10,11^. In the second, a spindle pushing force is mediated by the cytoplasmic F-actin meshwork ^12^. Both models enforce the notion that an F-actin-mediated force on the spindle, whether pushing or bulling, acts towards the nearest cortical side through which the PB is extruded. On the other hand, spindle orientation seems to be dependent on both F-actin and MTs ^16^. Astral-like MTs are only able to reach the cortex when the spindle is in very close proximity to the cortex ^16^. Because astral-like MTs are relatively short and cannot easily reach the cortex, they can only establish contacts with MTs nucleated by mcMTOCs which, in turn, could act as amplifying sites that anchor the spindle to the cortex. This model, which we here propose, thus depends on the presence of two opposing forces: mcMTOC-mediated MTs at one side and F-actin at the other side of the spindle (Fig. 7). These opposing forces would be essential to position the spindle centrally during pro-Met I and to prevent premature spindle migration, a risk factor for aneuploidy ^5^. Our model is consistent with three sets of observations: 1) mcMTOCs are exclusively localized asymmetrically, opposite the site of PB extrusion (the side of F-actin enrichment); 2) mcMTOCs undergo a significant decrease in number and volume during late Met I and Ana I/Telo I, allowing the F-actin mediated force to extrude the PB; and 3) nocodazole-mediated MT depolymerization advances the timing of chromosome migration to the cortex, which takes place at a relatively higher speed ^12,18^ whereas increasing mcMTOC numbers delays chromosome migration to the cortex, which occurs at a relatively reduced speed compared to controls.

**Figure 7:**
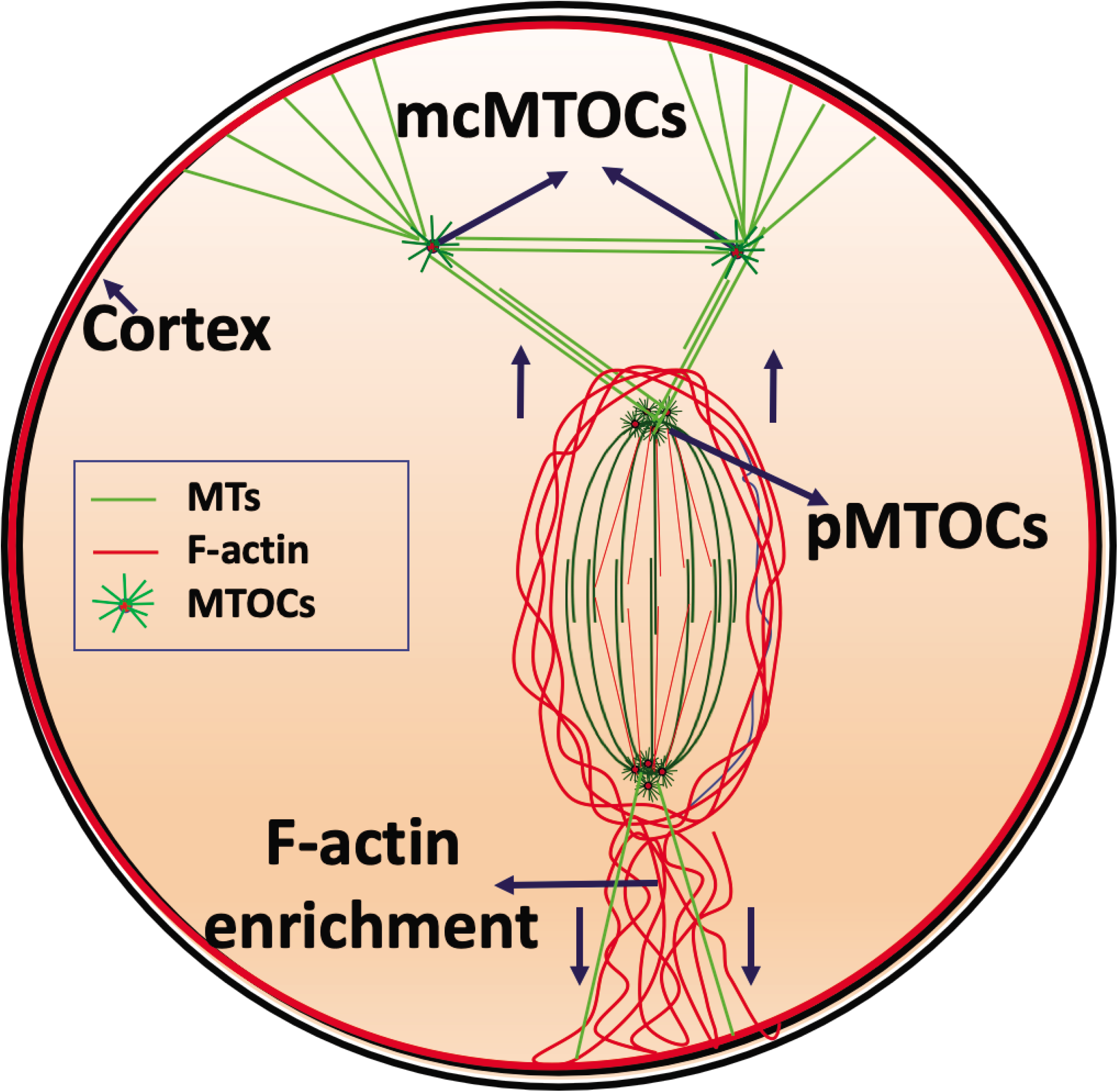
Schematic model for spindle positioning in the meiotic oocyte.

Depletion of mcMTOCs, disruption of mcMTOC numbers and MT depolymerization were each associated with abnormal spindle positioning and/or perturbed chromosome/spindle migration. For example, increasing mcMTOC numbers by 3-MA treatment significantly decreased chromosome speed and delayed chromosome migration towards the cortex. This phenotype accords with increased mcMTOC-mediated MT forces that oppose F-actin; thereby preventing proper spindle migration. Such phenotype was rescued, at least in part, by reducing mcMTOC numbers to the physiological range (6 to 8). Conversely, depleting mcMTOCs using laser ablation or MT depolymerization has the reciprocal effect ^12,18^. These findings indicate that the numbers of mcMTOCs must be regulated tightly to regulate spindle positioning and timely spindle migration.

In almost all mammals, including humans ^44,45^, meiotic oocytes contain numerous acentriolar MTOCs. This is in contrast to somatic mitotic cells, which contain only a pair of centrosomes that are sufficient to assemble and position the spindle centrally. Positioning the spindle at the center of mitotic cells depends on nucleating symmetrical astral MTs that anchor the spindle to the cell cortex. However, the mechanism appears different during MI. Mammalian eggs are large and the astral MTs from acentriolar MTOCs are relatively short. Yet, the cell must divide asymmetrically, something that would likely be difficult for a pair of symmetrical centrosomes to achieve. Our proposed model may, therefore, account for why meiotic oocytes rely upon two different functional sets of numerous MTOCs rather than a pair of typical centrosomes.

## MATERIAL AND METHODS

### Ethics

All animals were kept and experiments were conducted in accordance with the University of Missouri (Animal Care Quality Assurance Ref. Number, 9695) and UK Home Office regulations.

### Mouse strains

*Cep192-eGfp* reporter mice were generated by integrating a construct harboring the EGFP reporter gene into CF-1 mouse genome via CRISPR/Cas9-mediated homology-directed repair. The EGFP reporter was fused at the C-terminus of the endogenous mouse *Cep192* (Supplementary Fig. 3A). Homozygous breeding pairs of *Cep192-eGfp* reporter mice were used to maintain the colony.

### Oocyte collection, microinjection and culture

Full-grown GV-arrested oocytes were isolated from CF-1, C57BL/6 or *Cep192-eGfp* reporter CF-1 female mice (6-8-week-old) previously primed (∼44 h before collection), with pregnant mare serum gonadotropin (Lee BioSolutions #493-10-10) according to ^46,47^. Unless otherwise specified, CF-1 mice were used to conduct the experiments. Cumulus oocyte complexes (COCs) were collected and denuded using mechanical pipetting in bicarbonate-free minimal essential medium (MEM) containing 3 mg/ml polyvinylpyrolidone (PVP) and 25 mM Hepes (pH 7.3) supplemented with 2.5 μM milrinone (MilliporeSigma, St. Louis, MO, USA # M4659), a phospho diesterase inhibitor to arrest the oocytes at prophase I ^48^. Prophase I-arrested oocytes were microinjected with 10-15 pl of cRNAs encoding fluorescently labeled proteins while cultured in milrinone-containing MEM medium. Microinjected oocytes were then cultured in Chatot, Ziomek, and Bavister (CZB) medium ^49^ supplemented with milrinone in a humidified incubator with 5% CO_2_ in air at 37°C for ∼3 h to allow protein expression before releasing into milrinone-free CZB medium and initiating *in vitro* maturation. Met I, Ana I/Telo I or Met II oocytes were collected at 4, 7 or 14 h after NEBD.

Nocodazole (MilliporeSigma #M1404), MG-132 (MilliporeSigma #474790), Rapamycin (Enzo Life Sciences, Farmingdale, NY, USA #BML-A275), 3-Methyladenine (3-MA, Cayman Chemical, Ann Arbor, MI, USA #13242) were dissolved in dimethyl sulfoxide (DMSO) and used at a final concentration of 7.5 µM, 20 nM and 10 mM, respectively. *In vitro* maturation was carried out in organ culture dishes under humidified conditions (Becton Dickinson #353037).

### Cloning and *in vitro* cRNA synthesis

Generation of *Aurka-Gfp, H2b-mCherry* and *eGfp-Eb3* were described previously ^29,50^. DNA linearization of *Aurka-Gfp* and *H2b-mCherry* constructs was carried out using Nde I (New England BioLabs), whereas DNA linearization of *eGfp-Eb3* construct was carried out using SfiI (New England BioLabs). Purification of linearized DNA was carried out according to the manufacturer’s protocol (Qiagen, QIAquick PCR Purification). Purified DNA was *in vitro* transcribed using an mMessage mMachine T7 kit (Ambion) to generate *Aurka-Gfp* and *H2b-mCherry* cRNAs or mMessage mMachine T3 kit (Ambion) to generate *eGfp-Eb3* cRNA according to the manufacturer’s instructions. cRNA purification was performed using an RNAEasy kit (Qiagen) and stored at − 80°C.

### Immunocytochemistry and fluorescence microscopy

Meiotic oocytes were fixed for 20 min at room temperature in freshly prepared 2 % paraformaldehyde solution (MilliporeSigma #P6148) dissolved in phosphate buffer saline (PBS). Fixed oocytes were permeabilized in 0.1% Triton X-100 in PBS for 20 min prior to incubation for an additional 20 min in PBS containing 0.3% BSA and 0.01% Tween-20 (blocking solution). Primary antibody incubation was performed at room temperature for 1 h. Oocytes were then washed three times (8-9 min each) prior to incubation with secondary antibodies for 1 h. Oocytes were washed again three times in blocking solutions for 8-9 min. To detect F-actin, phalloidin (Texas Red X Phalloidin, ThermoFisher Scientific #T7471; 1:50) was added to secondary antibody solutions. Oocytes were mounted on slides using Vectashieled with 4’,6-Diamidino-2-Phenylindole, Dihydrochloride (DAPI; Vector Laboratories, Burlingame, CA, USA) to stain DNA. To label DNA for STED super-resolution imaging, 5□mg/m Hoechst 33342 was used (Molecular Probes H3570). The following primary antibodies were used in immunofluorescence: α-tubulin-Alexa Fluor 488 conjugate (Life Technologies #322 588; 1:100), **CEP192** (Young In Frontier #AR07-PA0001; 1:100), γ-tubulin (Millipore-Sigma #T6557; 1:75), Pericentrin (BD Biosciences #611814; 1:100), CREST autoimmune serum (Antibodies Incorporated #15-234; 1:25). Omitting the primary antibody served as a negative control. Fluorescence signals were detected under a 63X objective using Leica TCS SP8 confocal microscope equipped with 3-color, 3-D STED super-resolution 3X system. Images were captured to span the entire oocyte at 3 or 5 µm Z-intervals (confocal microscopy) or 0.5 µm Z-intervals (STED super-resolution microscopy). All images were acquired using the same laser power when the intensity of fluorescence is quantified.

### Time-lapse confocal microscopy

Oocytes expressing fluorescently labeled proteins were transferred to milrinone-free CZB medium and imaged over time under a 63X objective using Leica TCS SP8 confocal microscope equipped with microenvironmental chamber to maintain the oocytes at controlled CO_2_ (5%) and temperature (37 °C) in a humidified air. DIC, GFP and mCherry image acquisitions were started at prophase I stage or as indicated and images were captured every certain time according to each experimental design (as indicated in corresponding figure legends). Images were captured to span the entire oocyte at 3 or 5 µm Z-intervals.

### Depletion of mcMTOCs using laser ablation

Depletion of mcMTOCs was carried out using two-photon laser ablation which has the advantage of offering deeper tissue penetration, efficient ablation and minimizing off-target effects ^39^. Two different microscopes were used. Oocytes expressing fluorescently labeled proteins were transferred to milrinone-free MEM medium and mcMTOCs were ablated using a laser with 925 nm wavelength of upright LaVision BioTec TriM Scope II (with controlled temperature at 37 °C) or to milrinone-free CZB medium and mcMTOCs were ablated using a laser with 820 nm wavelength if Leica TCP SP8 two-photon inverted microscope (equipped with microenvironmantal chamber to control CO_2_ and temperature) was used. In both cases, a small square area(s) surrounding mcMTOCs were marked and then exposed to a laser beam. We compared the first image (before ablation) and the second image (after ablation) in the time series cycle to ensure that after ablation, the fluorescence in the targeted mcMTOCs decreased to that observed at the background levels. Next, we moved the focal plane and ablated the remaining mcMTOCs. McMTOC-depleted oocytes underwent live imaging using the same parameters. Control oocytes were exposed to the same protocol except ablating random areas of the cytoplasm, just adjacent and equal to the same size and number of mcMTOCs.

### Image processing and analysis

Images acquired using 3-D STED super-resolution microscopy were deconvolved using Huygens Professional software before image analysis. NIH image J software (National Institute of Health, Bethesda, MD, USA) was used to process and analyze the images of fixed oocytes. The speed and average distance of chromosome/spindle/MTOCs at their final position over time were analyzed using the manual tracking function of NIH image J software. The point of intersection between the line connecting the two dominant spindle poles and spindle midzone was used to determine the position of the spindle. The point of intersection between the two lines representing the minor axis length and the major axis length of chromosomes was used to determine the position of all chromosomes ^51^. The speed and distance for each MTOC were analyzed separately before calculating the average for all MTOCs within each oocyte. 3D reconstruction of MTOCs, MTOC number and volume were processed and analyzed using isosurface spot analysis feature of Imaris software (Bitplane, Zürich, Switzerland) according to ^19^. Briefly, based on MTOC signal to noise, the threshold value was adjusted on an oocyte-to-oocyte basis followed by MTOC surface segmentation. MTOC number was calculated using the spot analysis feature, whereas MTOC volume was analyzed using surface analysis feature. mcMTOCs were quantified after excluding the pMTOCs manually, and *vice versa*. Same processing parameters were applied for each experimental analysis. Spindle orientation was analyzed using Imaris measurement points. The two dominant spindle poles were determined automatically based on increased MT fluorescent intensity at spindle poles. A line connecting the two spindle poles defined the spindle axis at each time point. The angle between two successive spindle axes was computed to determine the spindle orientation angle.

### *In situ* chromosome counting

Oocytes at Met II stage (12 h post-NEBD) were treated with 100 µM monastrol (MilliporeSigma #M8515), an Eg5-kinesin inhibitor to induce monopolar spindle formation with subsequent chromosome dispersion ^40,41^. Oocytes were fixed and immunostained as previously mentioned with CREST autoimmune serum to detect kinetochores. Oocytes were then mounted onto a glass slide using Vectashield with DAPI (Vector Laboratories) to label DNA. Confocal microscopy was used to image the entire region of the chromosomes at 0.7-µm Z-intervals to capture all kinetochores. Serial confocal sections were analyzed and the total number of kinetochores were counted using NIH image J software.

### Statistical analysis

One-way ANOVA, Student t-test and chi-square contingency test were used to evaluate the differences between groups using GraphPad Prism. ANOVA test was followed by the Tukey post hoc test to allow the comparison among groups. The differences of P < 0.05 were considered significant. The data were expressed as means ± SEM.

## ACKNOWLEDGMENTS

The authors would like to thank Dr. David Glover, California Institute of Technology, USA for critical reading and editing the manuscript, and for providing resources. The authors would like to thank all members of the Glover lab and the Balboula lab for valuable help and discussions.

## FIGURE LEGENDS

**Supplementary Figure 1:**
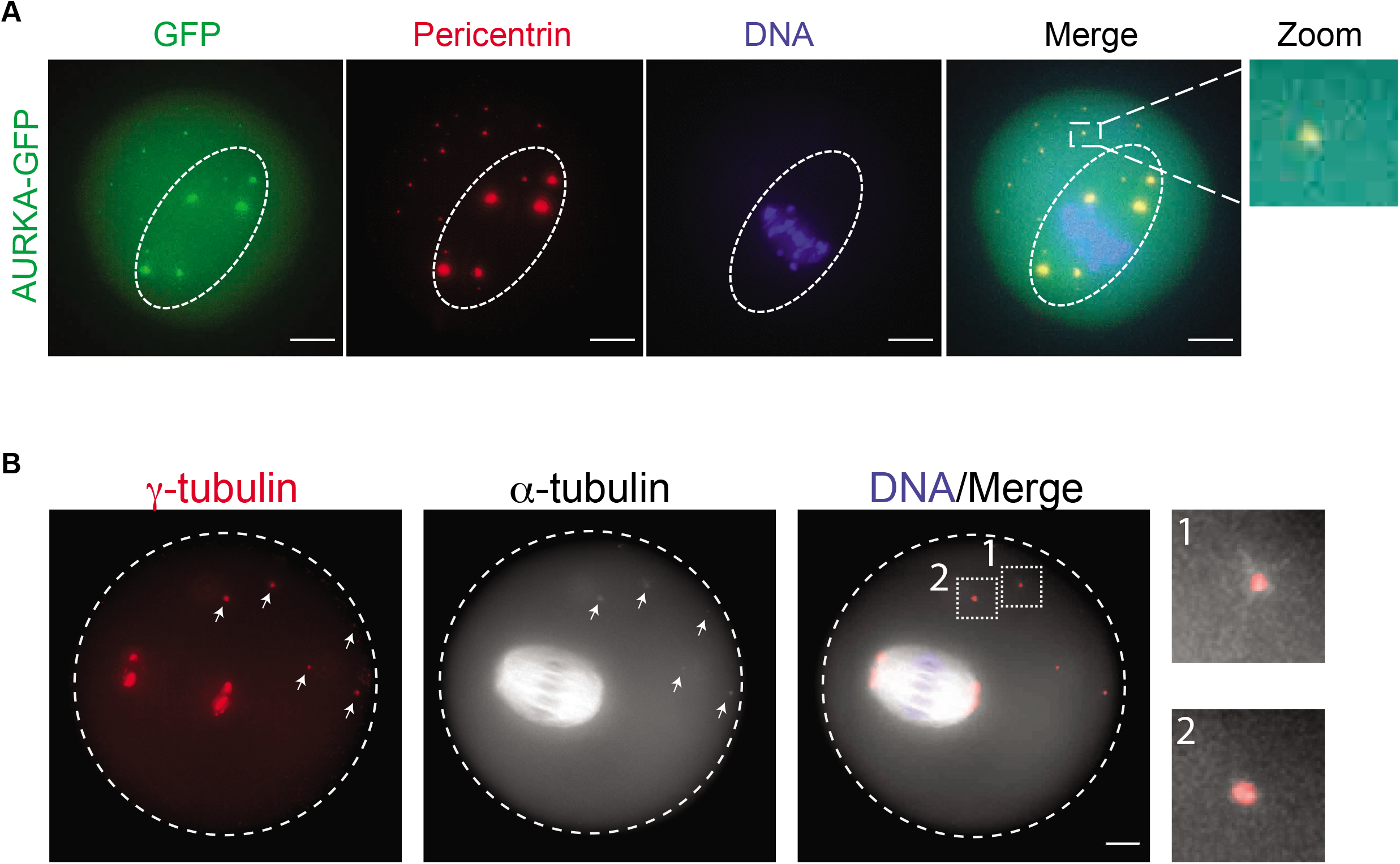
**(A) AURKA-GFP foci colocalize with pericentriolar material protein (pericentrin) at mcMTOCs.** Full-grown prophase-I oocytes were injected with *Aurka-Gfp* (green), incubated in milrinone-containing CZB medium for 3 h, followed by *in vitro* maturation. Metaphase I oocytes were fixed and immunostained using pericentrin antibody to label MTOCs (red). DAPI was used to detect DNA (blue). (B) **MTOCs in the cytoplasm and at the spindle poles of meiotic oocytes collected from C57BL/6 mice**. Full-grown prophase-I oocytes collected from C57BL/6 mice were *in vitro* matured for 7 h. Metaphase I oocytes were fixed and immunostained using γ-tubulin and α-tubulin antibodies to label MTOCs (red) and microtubules (pseudo grey). DAPI was used to detect DNA (blue). White arrows represent mcMTOCs. Shown are representative Z-projection of confocal images. Scale bars represent 10□μm.

**Supplementary Figure 2:**
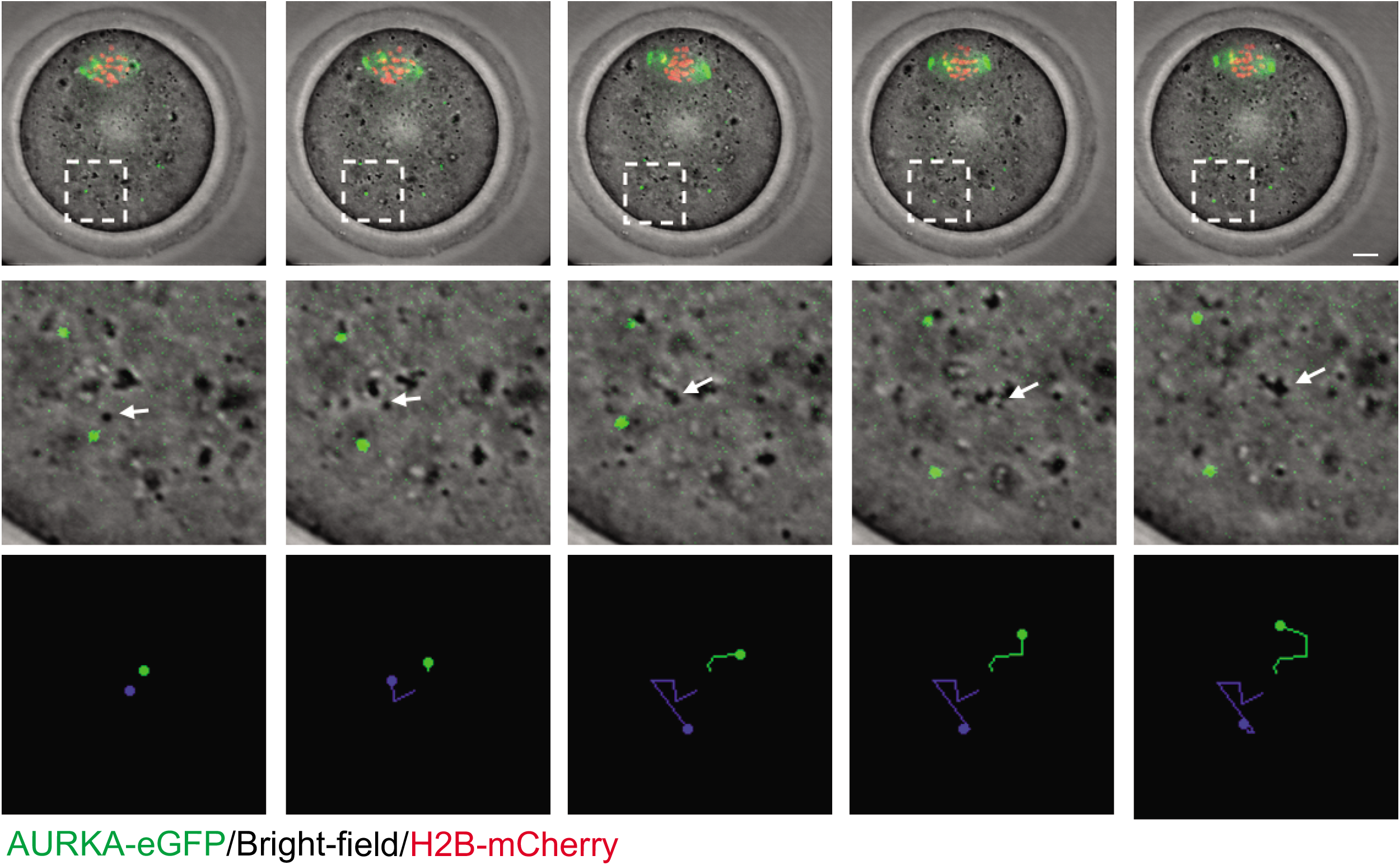
mcMTOC movement does not follow the movement of the cytoplasm. Representative images (Z-projection of 16 sections every 3□μm) of time-lapse confocal microscopy of live oocytes expressing AURKA-GFP (MTOCs, green) and H2B-mCherry (chromosomes) from a time course. Fluorescence and bright-field images were captured every 15 min (time, h:min). The white arrow represents a cytoplasmic droplet. Lower panels show the tracking of both the cytoplasmic droplet (green) and an mcMTOC (blue). The scale bar represents 10□μm.

**Supplementary Figure 3:**
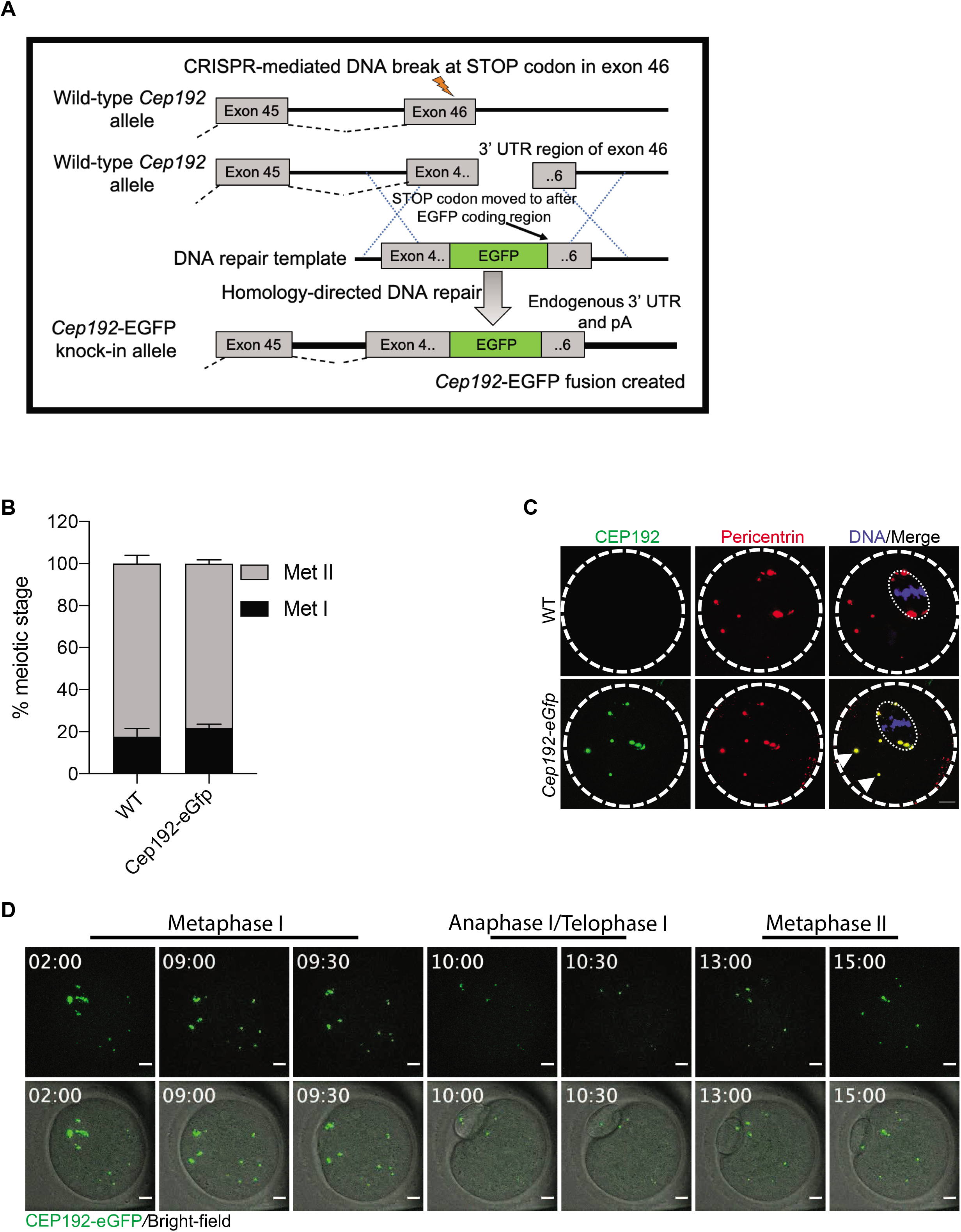
(A) Diagram for generating *Cep192-eGfp* reporter mice. (B) Full-grown prophase I-arrested oocytes were collected from wild-type (WT) and *Cep192-eGfp* reporter mice, *in vitro* matured for 16 h (Met II) and assessed for meiotic progression. (C) Full-grown prophase I-arrested oocytes were collected from *Cep192-eGfp* reporter mice (CEP192 MTOCs are labeled green) and *in vitro* matured for 7 h. Metaphase I oocytes were fixed and immunostained using pericentrin antibody to label MTOCs (red). DAPI was used to detect DNA (blue). Arrowheads represent colocalized CEP192 with pericentrin at mcMTOCs. Shown are representative Z-projection images. Scale bars represent 10□μm. (D) Representative images (Z-projection of 13 sections every 5□μm) of time-lapse confocal microscopy of a live *Cep192-eGfp* oocyte from a time course (see Supplemental Movie 7). Fluorescence and bright-field images (lower panels) were captured every 30 min post NEBD (time, h:min). Scale bars represent 10□μm.

**Supplementary Figure 4:**
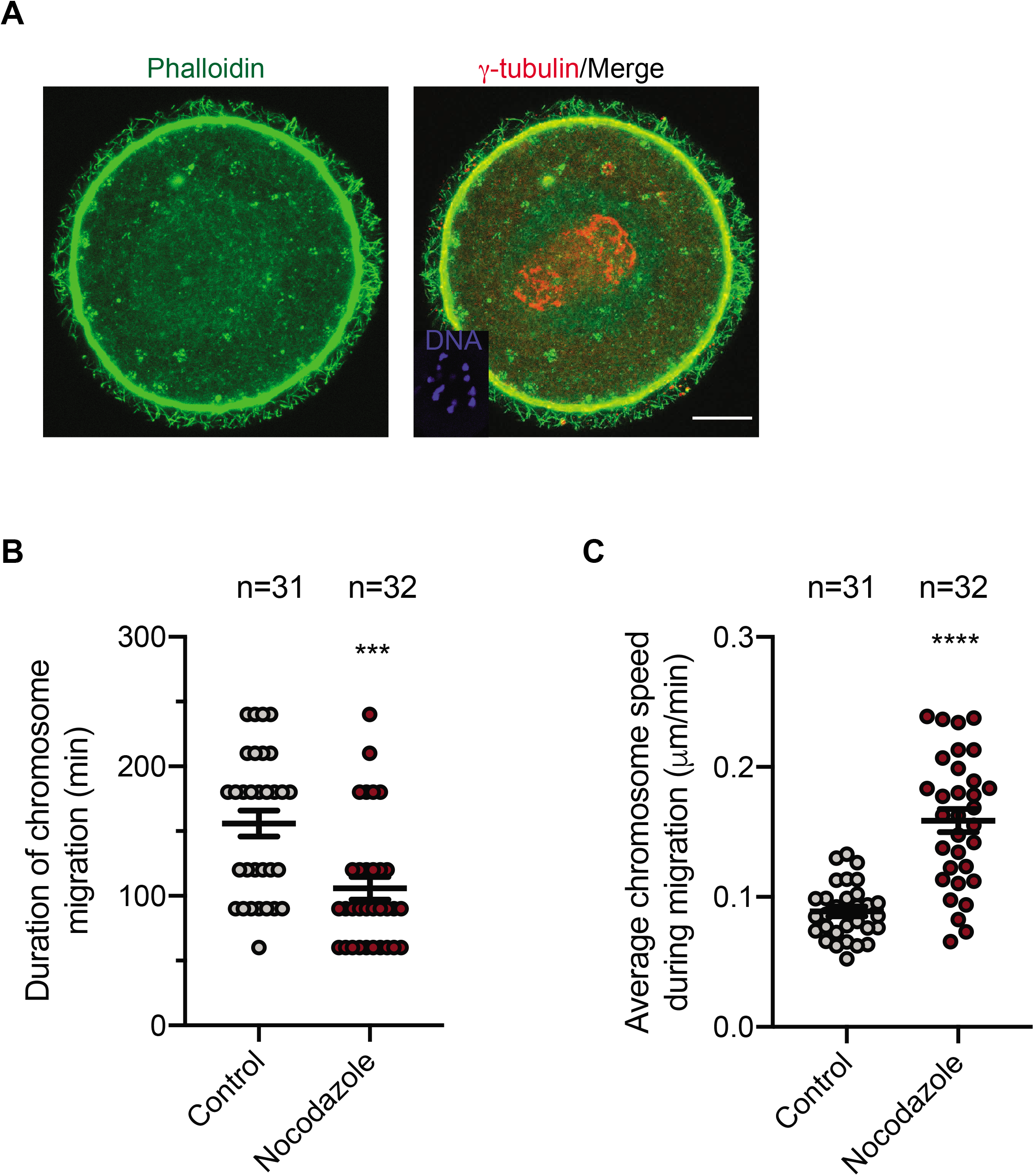
**(A) F-actin localization in mouse oocytes.** Full-grown prophase-I oocytes were *in vitro* matured for 7 h. Metaphase I oocytes were fixed and immunostained using γ-tubulin to label MTOCs (red). Hoechst stain was used to detect DNA (blue) and phalloidin stain was used to detect F-actin (pseudo green). Shown are representative Z-projection. The scale bar represents 10□μm. **(B**,**C) Inhibition of MTs accelerates chromosome migration towards the cortex in meiotic oocytes**. Full-grown prophase-I oocytes were divided into two groups and treated with DMSO or nocodazole (added at 0 h after collection) followed by *in vitro* maturation and time-lapse imaging. Images were captured every 15 min. (B) Quantification of the average time spent by chromosomes until reaching the cortex. (C) Quantification of average chromosome speed during migration. The data are expressed as mean ± SEM. Student-t test was used to analyze the data. Values with asterisks vary significantly, ***P < 0.001, ****P < 0.0001. The total number of analyzed oocytes (from two independent replicates) is specified above each graph.

**Supplementary Figure 5:**
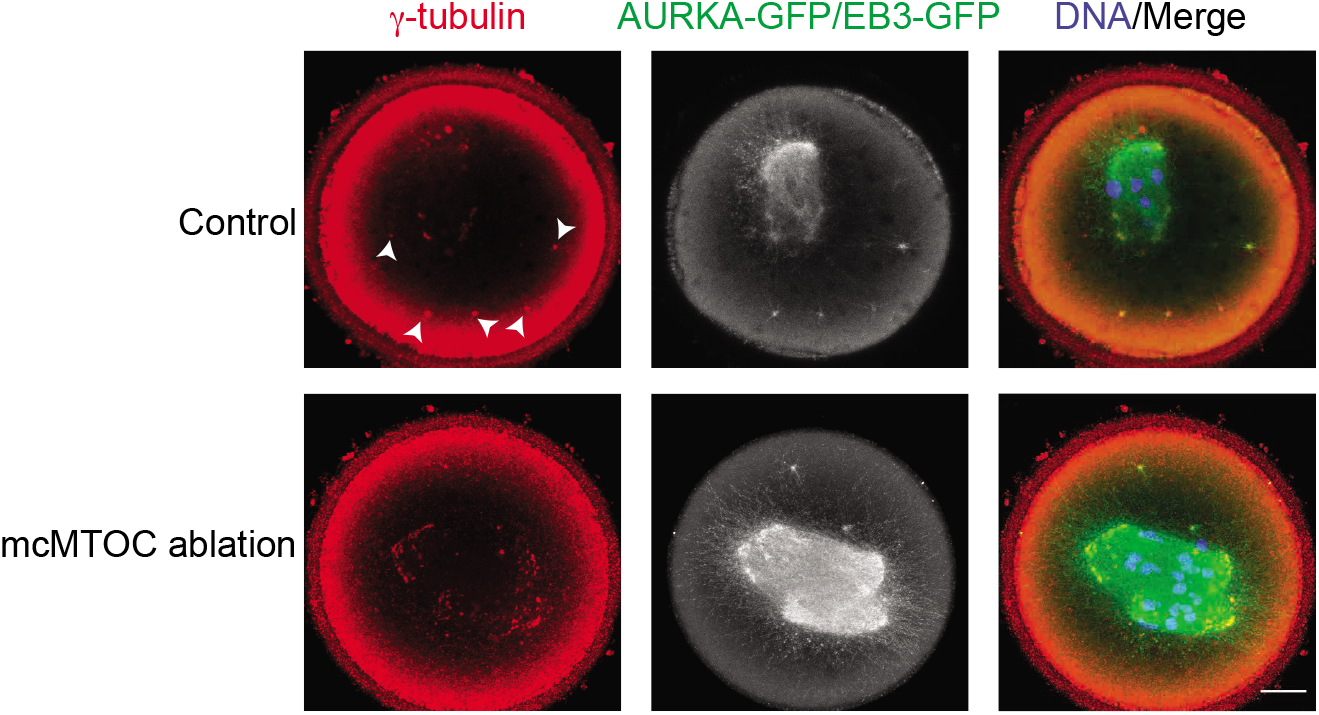
Two-photon laser ablation efficiently depletes mcMTOCs. Oocytes expressing AURKA-GFP and eGFP-EB3 were *in vitro* maturated for 6 h (metaphase I, Met I), transferred to CZB medium with MG-132, followed by mcMTOC depletion using two-photon laser ablation. Small square area(s) surrounding mcMTOCs were marked and then exposed to a laser with 820 nm wavelength. Control oocytes were exposed to the same parameters except ablating random areas of the cytoplasm, just adjacent and equal to the same size and number of mcMTOCs. Control and mcMTOC-ablated oocytes were fixed and immunostained using γ-tubulin antibody to label MTOCs. DAPI was used to detect DNA (blue). Arrowheads represent mcMTOCs. Shown are representative Z-projection of confocal images. The scale bar represents 10□μm.

**Supplementary Figure 6:**
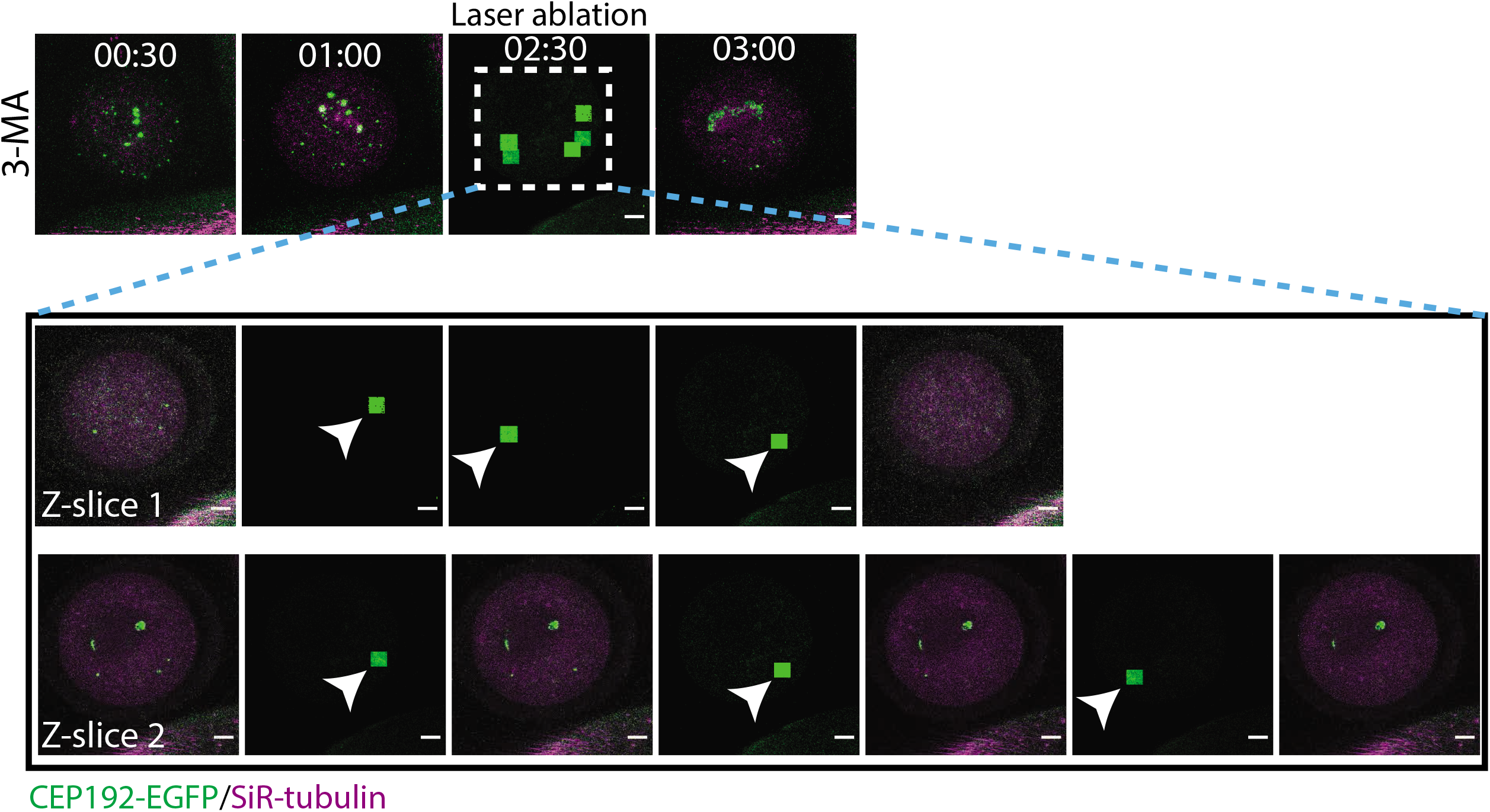
Representative Z-sections of 3D time-lapse imaging of a 3-MA-treated oocyte during mcMTOC ablation. Full-grown prophase I-arrested oocytes were collected from *Cep192-eGfp* reporter mice (MTOCs are labeled green) and *in vitro* matured in CZB medium containing SiR-tubulin (to label microtubules, magenta) for 2 h. 3-MA was added to the maturation medium at NEBD. At prometaphase I, 6 out of 14 mcMTOCs were depleted using laser ablation. Small square area(s) surrounding mcMTOCs were marked and then exposed to a laser with 820 nm wavelength. White arrowheads represent laser beam targets. Shown are the two Z-sections in which 6 mcMTOCs were ablated. Same oocyte as shown in Fig. 6F (lower panels). The scale bar represents 10□μm.

**Supplementary Movie 1:** Time-lapse confocal microscopy of MTOCs in live oocyte. The full-grown prophase-I oocyte was injected with cRNAs encoding *H2b-mCherry* (red) and *Aurka-Gfp* (pseudo grey), followed by *in vitro* maturation. Fluorescence images (Z-projection of 16 sections every 3□μm) were captured every 20 min (time, h:min). Same oocyte as shown in Fig. 1A (upper panels). The scale bar represents 10□μm.

**Supplementary Movie 2:** 3D reconstruction of MTOCs from the oocyte in Supplementary Movie 1. Same oocyte as shown in Fig. 1B (lower panels).

**Supplementary Movie 3:** 3D reconstruction of MTOCs from the oocyte in Supplementary Movie 1 during metaphase I.

**Supplementary Movie 4:** Time-lapse confocal microscopy of MTOCs in live oocyte. The full-grown prophase-I oocyte was injected with cRNAs encoding *H2b-mCherry* (red) and *Aurka-Gfp* (pseudo grey), followed by *in vitro* maturation. Fluorescence images (Z-projection of 16 sections every 3□μm) were captured every 15 min (time, h:min). The scale bar represents 20□μm.

**Supplementary Movie 5:** Tracking of 3D reconstructed mcMTOCs from the oocyte in Supplementary Movie 4 during nuclear envelope breakdown (NEBD) to early metaphase I (Met I). Same oocyte as shown in Fig. 2A.

**Supplementary Movie 6:** Tracking of 3D reconstructed mcMTOCs from the oocyte in Supplementary Movie 4 during late metaphase I to telophase I. Same oocyte as shown in Fig. 2C.

**Supplementary Movie 7:** Time-lapse confocal microscopy of MTOCs in live oocyte. Full-grown prophase I-arrested oocytes were collected from *Cep192-eGfp* reporter mice (MTOCs are labeled green) and *in vitro* matured in CZB medium. Fluorescence and bright-field images (Z-projection of 13 sections every 5□μm) were captured every 30 min (time post NEBD, h:min). The scale bar represents 10□μm.

**Supplementary Movie 8:** Time-lapse confocal microscopy of MTOCs in live oocyte. The full-grown prophase-I oocyte was injected with cRNAs encoding *H2b-mCherry* (red) and *Aurka-Gfp* (pseudo grey), followed by *in vitro* maturation. Fluorescence and bright-field images (Z-projection of 16 sections every 3□μm) were captured every 20 min (time, h:min). Same oocyte as shown in Fig. 3C (lower panels). The scale bar represents 10□μm.

**Supplementary Movie 9:** 3D time-lapse imaging of live oocyte expressing AURKA-GFP (pseudo grey) and eGFP-EB3 (pseudo grey) during mcMTOC ablation. The white squares represent the actual laser beam targets. Same oocyte as shown in Fig. 4B.

**Supplementary Movie 10:** Tracking the spindle over time in control oocyte (exposed to ablating random areas of the cytoplasm) expressing AURKA-GFP (pseudo grey) and eGFP-EB3 (pseudo grey), while cultured in MG-132-containing medium for 9 h, using 3D time-lapse microscopy. Fluorescence images were captured every 3 min (time post laser ablation, h:min). The scale bar represents 10□μm. Same oocyte as shown in Fig. 4E.

**Supplementary Movie 11:** Tracking the spindle over time in mcMTOC-laser-ablated oocyte expressing AURKA-GFP (pseudo grey) and eGFP-EB3 (pseudo grey), while cultured in MG-132-containing medium for 9 h, using 3D time-lapse microscopy. Fluorescence images were captured every 3 min (time post laser ablation, h:min). The scale bar represents 10□μm. Same oocyte as shown in Fig. 4E.

**Supplementary Movie 12:** Tracking the spindle over time in control oocyte during meiosis I. Full-grown prophase I-arrested oocytes were collected from *Cep192-eGfp* reporter mice (MTOCs are labeled green) and *in vitro* matured in CZB medium containing SiR-tubulin (to label microtubules, magenta) until Met I prior to control laser ablation (oocytes were exposed to ablating multiple areas of the cytoplasm but not mcMTOCs). Fluorescence and bright-field images (Z-projection of 18 sections every 3□μm) were captured every 30 min (time post laser ablation, h:min). Same oocyte as shown in Fig. 4G (upper panels). The scale bar represents 10□μm.

**Supplementary Movie 13:** Tracking the spindle over time in mcMTOC-ablated oocyte during meiosis I. Full-grown prophase I-arrested oocytes were collected from *Cep192-eGfp* reporter mice (MTOCs are labeled green) and *in vitro* matured in CZB medium containing SiR-tubulin (to label microtubules, magenta) until Met I. mcMTOCs were depleted using laser ablation prior to time-lapse confocal microscopy. Fluorescence and bright-field images (Z-projection of 18 sections every 3□μm) were captured every 30 min (time post laser ablation, h:min). Same oocyte as shown in Fig. 4G (middle panels). The scale bar represents 10□μm.

**Supplementary Movie 14:** Tracking the spindle over time in another example of mcMTOC-ablated oocyte during meiosis I. Full-grown prophase I-arrested oocytes were collected from *Cep192-eGfp* reporter mice (MTOCs are labeled green) and *in vitro* matured in CZB medium containing SiR-tubulin (to label microtubules, magenta) until Met I. mcMTOCs were depleted using laser ablation prior to time-lapse confocal microscopy. Fluorescence and bright-field images (Z-projection of 18 sections every 3□μm) were captured every 30 min (time post laser ablation, h:min). Same oocyte as shown in Fig. 4G (lower panels). The scale bar represents 10□μm.

**Supplementary Movie 15:** Tracking the chromosomes over time in DMSO-treated oocytes during meiosis I. Bright-field images (Z-projection of 16 sections every 3□μm) were captured every 30 min (time, h:min). NEBD represents 0 h. Same oocyte as shown in Fig. 6A (Control). The scale bar represents 10□μm.

**Supplementary Movie 16:** Tracking the chromosomes over time in Rapamycin-treated oocytes during meiosis I. Bright-field images (Z-projection of 16 sections every 3□μm) were captured every 30 min (time, h:min). NEBD represents 0 h. Same oocyte as shown in Fig. 6A (Rapamycin). The scale bar represents 10□μm.

**Supplementary Movie 17:** Tracking the chromosomes over time in 3-MA-treated oocytes during meiosis I. Bright-field images (Z-projection of 16 sections every 3□μm) were captured every 30 min (time, h:min). NEBD represents 0 h. Same oocyte as shown in Fig. 6A (3-MA, upper panels). The scale bar represents 10□μm.

**Supplementary Movie 18:** Tracking the chromosomes over time in 3-MA-treated oocytes during meiosis I (another representative showing defective asymmetrical division). Bright-field images (Z-projection of 16 sections every 3□μm) were captured every 30 min (time, h:min). NEBD represents 0 h. Same oocyte as shown in Fig. 6A (3-MA, lower panels). The scale bar represents 10□μm.

**Supplementary Movie 19:** Tracking the spindle over time in DMSO-treated oocytes during meiosis I. Full-grown prophase I-arrested oocytes were collected from *Cep192-eGfp* reporter mice (MTOCs are labeled green) and *in vitro* matured in CZB medium containing SiR-tubulin (to label microtubules, magenta). Time-lapse imaging started at prometaphase I (3 h post NEBD). Fluorescence images (Z-projection of 13 sections every 5□μm) were captured every 45 min (time, h:min). Same oocyte as shown in Fig. 6F (upper panels). The scale bar represents 10□μm.

**Supplementary Movie 20:** Tracking the spindle over time in 3-MA-treated oocytes during meiosis I. Full-grown prophase I-arrested oocytes were collected from *Cep192-eGfp* reporter mice (MTOCs are labeled green) and *in vitro* matured in CZB medium containing SiR-tubulin (to label microtubules, magenta) for 2 h. 3-MA was added to the maturation medium at NEBD. Time-lapse imaging started at prometaphase I (3h post NEBD). Fluorescence images (Z-projection of 13 sections every 5□μm) were captured every 45 min (time, h:min). Same oocyte as shown in Fig. 6F (middle panels). The scale bar represents 10□μm.

**Supplementary Movie 21:** Tracking of 3D reconstructed spindle poles from the oocyte in Supplementary Movie 20. Same oocyte as shown in Fig. 6F (middle panels).

**Supplementary Movie 22:** Tracking the spindle over time in 3-MA-treated oocytes exposed to partial mcMTOC laser ablation during meiosis I. Full-grown prophase I-arrested oocytes were collected from *Cep192-eGfp* reporter mice (MTOCs are labeled green) and *in vitro* matured in CZB medium containing SiR-tubulin (to label microtubules, magenta) for 2 h. 3-MA was added to the maturation medium at NEBD. When the oocyte reached prometaphase I, 6 out of 14 mcMTOCs were depleted using laser ablation (Supplementary Fig. 6). Time-lapse imaging started 1h after laser ablation. Fluorescence images (Z-projection of 13 sections every 5□μm) were captured every 45 min (time, h:min). Same oocyte as shown in Fig. 6F (lower panels). The scale bar represents 10□μm.

